# A competitive activity-based protein profiling platform yields cell wall synthesis inhibitors active against replicating and non-replicating *Mycobacterium tuberculosis*

**DOI:** 10.1101/2021.04.16.440156

**Authors:** Michael Li, Hiren V. Patel, Armand B. Cognetta, Trever C. Smith, Ivy Mallick, Jean-François Cavalier, Stephane Canaan, Bree B. Aldridge, Benjamin F. Cravatt, Jessica C. Seeliger

## Abstract

The identification and validation of a small molecule’s targets is a major bottleneck in the discovery process for tuberculosis antibiotics. Activity-based protein profiling (ABPP) is an efficient tool for determining a small molecule’s targets within complex proteomes. However, how target inhibition relates to biological activity is often left unexplored. Here we studied the effects of 1,2,3-triazole ureas on *Mycobacterium tuberculosis* (*Mtb*). After screening ~200 compounds, we focused on two inhibitors active against both exponentially replicating and hypoxia-induced drug-tolerant *Mtb* that form part of a four-compound structure-activity series. The compound with negligible activity revealed potential false positive targets not addressed in other ABPP studies. Biochemistry, computational docking, and morphological analysis confirmed that active compounds preferentially inhibit serine hydrolases with cell wall and lipid metabolism functions and that disruption of the cell wall underlies biological activity. Our findings showed that ABPP identifies the targets most likely relevant to a compound’s antibacterial activity.

## INTRODUCTION

The rising incidence of antibiotic resistance in the causative bacterium *Mycobacterium tuberculosis* (*Mtb*) makes the need to develop novel TB therapies ever more urgent. *Mtb* inhibitor discovery has relied primarily on two approaches: phenotypic compound screening or target-based compound screening. In both cases, an important step for compound optimization is the identification and validation of a compound’s target. Target validation has depended largely on high-throughput genetic methods such as generating spontaneous genomic mutations (Andries et al., 2005; Christophe et al., 2009; Grzegorzewicz et al., 2012; Makarov et al., 2009; Pethe et al., 2013; Remuiñán et al., 2013; Stanley et al., 2013; Stover et al., 2000) and overexpressing or underexpressing putative targets(Abrahams et al., 2012; Evans and Mizrahi, 2015; Johnson et al., 2019; Krieger et al., 2012; Wei et al., 2011). These genetic methods are most useful when an inhibitor acts predominantly via a single target or pathway, but less revealing when multiple targets underlie biological activity.

Indeed, the ability to act on multiple targets is an important feature of some antibiotics, including the first-line tuberculosis drug isoniazid(Argyrou et al., 2006b, 2006a; Gangadharam et al., 1963; Silver, 2007). However, spontaneous mutations that confer resistance can reveal the most easily mutated targets to a compound, but not necessarily all the targets relevant to compound activity. There is thus a need for a methodology that detects all potential targets simultaneously and thus provides a comprehensive and accurate assessment of an inhibitor’s mechanism of action.

To achieve this goal, activity-based protein profiling (ABPP) has emerged as an efficient tool that can monitor the reactivity of nucleophiles within complex proteomes. In *Mtb*, as in other organisms, the broad utility of ABPP has inspired the modification of covalent inhibitors into probes, usually by addition of an alkyne for subsequent azide-alkyne cycloaddition to attach a tag to enrich and detect a compound’s targets of the compounds (Lehmann et al., 2016, 2018; Ravindran et al., 2014) or to investigate activity of particular enzymes (Duckworth et al., 2012; Lentz et al., 2016). The most common chemical modification is the addition of an alkyne that can be coupled by azide-alkyne cycloaddition to a tag for enrichment and identification.

An alternative to inhibitor modification is to use ABPP in a competitive format. Here, inhibitor and ABP target the same reactive nucleophile, such that the inhibitor is not expected to exert its biological effects via targets other than those detected by ABPP. In *Mtb*, competitive ABPP has been used to identify the targets of staurosporine, a non-covalent, ATP-competitive inhibitor. Staurosporine-treated cells were labeled with an ATP analog that was also used to annotate ATP-binding proteins in Mtb(Ansong et al., 2013). With this strategy PknF, PknD and PknB were identified as the ATP-hydrolyzing enzymes most strongly inhibited by staurosporine. Competitive ABPP has also been applied to identify the serine hydrolase targets of oxadiazolone compounds and the cyclipostin analog CyC_17_ (Nguyen et al., 2017, 2018a). With this strategy, enzymes related to cell wall biosynthesis and lipid metabolism were identified. However, the relative contribution of these targets to the biological activity of the inhibitors is not well understood, as overexpression of individual targets did not significantly alter mycobacterial susceptibility to CyC_17_ or oxadiazolone compounds.

In the present study we used competitive ABPP to identify the serine hydrolase targets of 1,2,3-triazole ureas that inhibit *Mtb* growth and to quantify the inhibition of selected purified serine hydrolases by these compounds. We have indeed previously reported that 1,2,3-triazole ureas are potent and selective inhibitors of serine hydrolases through covalent inhibition of the active site serine(Adibekian et al., 2011; Hsu et al., 2013b, 2013a). We reasoned that using competitive ABPP to discover the targets in *Mtb* would delineate key pharmacologically targetable serine hydrolases and their associated pathways. To prioritize targets, we used a four-compound structure-activity series to test the hypothesis that enzymes that are preferentially inhibited by active vs. inactive compounds are more likely contributing to the observed antibacterial activity of each inhibitor. Biochemical assays and computational docking validated the structure-activity relationships among the selected inhibitors and supported our use of ABPP to prioritize serine hydrolase targets. The functions of prioritized targets suggested activity via the inhibition of cell wall and lipid synthesis and we corroborated this proposed mechanism of action using morphological profiling of inhibitor-treated *Mtb*.

## EXPERIMENTAL PROCEDURES

### Phenotypic screening for inhibition of Mtb growth

Autoluminescent *Mtb* were grown to initial OD_600_ 0.5-0.6 and subcultured to OD_600_ 0.02 in modified Roisin’s medium. Where noted, 0.5% glycerol was substituted with 100 μg/mL cholesterol (from 500X stock in 1:1 (v/v) ethanol/Tyloxapol). The triazole urea library was obtained from the Cravatt laboratory (Adibekian et al., 2011). The final concentration of all compounds was 10 μM and the final concentration of DMSO was 1% (v/v). Plates were incubated in a humidified incubator at 37 °C with 5% CO_2_ for 7 days. Luminescence was measured with 500 ms integration time (Molecular Devices SpectraMax M3). Percent inhibition was calculated as 100 × [(μDMSO ‒ μCompound)/ μDMSO] where μ is the average autoluminescent signal.

### Target identification by ABPP-SILAC

For all experiments, *Mtb* was incubated with ~1X MIC AA691 or ~1X MIC AA692 and maintained at a moderate cell density to reflect conditions under which antibacterial activity was observed and to minimize clumping and promote even and reproducible contact between bacterial cells and inhibitors. *Mtb* was cultured to an OD_600_ ~1 in modified Roisin’s medium containing either ^14^N (“light”) or ^15^N (“heavy”) ammonium chloride as a sole nitrogen source. Light cultures were incubated with 2 μM AA691, AA692 or AA702 and heavy cultures were incubated with 0.02% (v/v) DMSO vehicle control for 2 h. The cell pellet was washed with 1 mL 0.05% Tween 20 in PBS and then 2 mL PBS. Cell suspensions were lysed by bead beating at 4000 rpm for 45 s on/off cycles (Benchmark Scientific BeadBug homogenizer). Protein concentration in clarified lysates was measured by the BCA assay (Pierce) and lysates were diluted to 2 mg/mL with PBS in 0.5 mL total volume. The lysates were treated with 10 μM fluorophosphonate-biotin (FP-biotin; gift of Dr. Eranthie Weerapana) for 1 h at 22 °C and desalted (PD Miditrap G-25 columns, GE Healthcare). Each light lysate was then combined 1:1 with a heavy lysate in 1 mL total volume and incubated with 2% SDS and 2 M urea final concentration in PBS in a final volume of 2 mL for 40 min at 22 °C, 110 rpm. The combined lysates were then spun at 4300 × *g* for 20 min. The supernatants were diluted with 8 mL PBS and filtered twice through a 0.2-μm polyethersulfone filter. At this point the filtered lysates were deemed non-viable based on viability testing for *Mtb* and removed from the Biosafety Level 3 laboratory for further analysis.

### MorphEUS analysis

*Mtb* was incubated for 21 h with AA691, AA692, AA701, and AA702 at 50 μM or 500 μM in 7H9 medium supplemented with OADC. These concentrations correspond to ~0.5X and 5X the MIC of AA692 under these conditions; the MIC of AA692 is higher in 7H9 (~100 μM) than in Roisin (11 μM; **Table 1**). The final concentration of DMSO was 1% (v/v). Compound-treated *Mtb* cultures were fixed with 4% paraformaldehyde (Alfa Aesar, 43368) for 1 h, washed twice with 100 μL PBS containing 0.2% Tween-80 (PBST), and resuspended in 100 μL PBST. Staining and imaging were as described previously (Smith et al., 2020). Briefly, 50 μL of fixed *Mtb* cells were diluted with 50 μL PBST and stained with 0.6 μg of FM4-64-FX (ThermoFisher, F34653) and 15 μl of 0.1 μM SYTO 24 (ThermoFisher, S7559) at room temperature in the dark for 30 min. Once stained, the cells were washed once with 100 μL of PBST and resuspended in 30 μL of PBST. Stained *Mtb* were spotted onto agarose plates (1% w/v agarose; SigmaAldrich A3643-25G) and images were captured with a widefield DeltaVision PersonalDV (Applied Precisions) microscope. Bacteria were illuminated using an InsightSSI Solid State Illumination system with transmitted light for phase contrast microscopy and a DV Elite CMOS camera. SYTO 24 was imaged using 475 nm excitation and 525 nm emission. FM4-64-FX was imaged with 475 nm excitation and 679 nm emission. Two technical replicate images were taken from each sample for a total of 50 images per biological replicate. Three biological replicates were generated for each drug treatment.

**Table 1.**
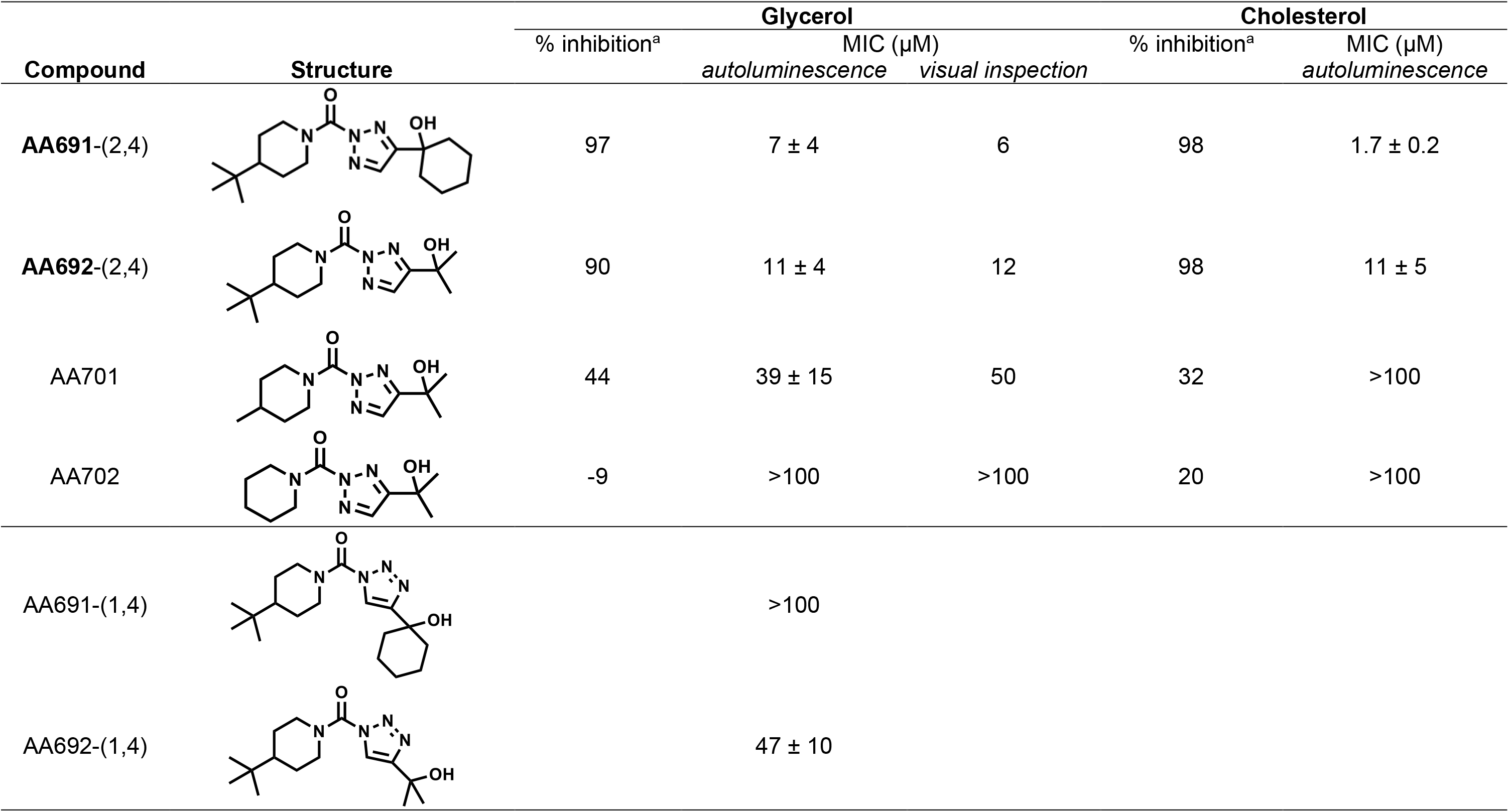
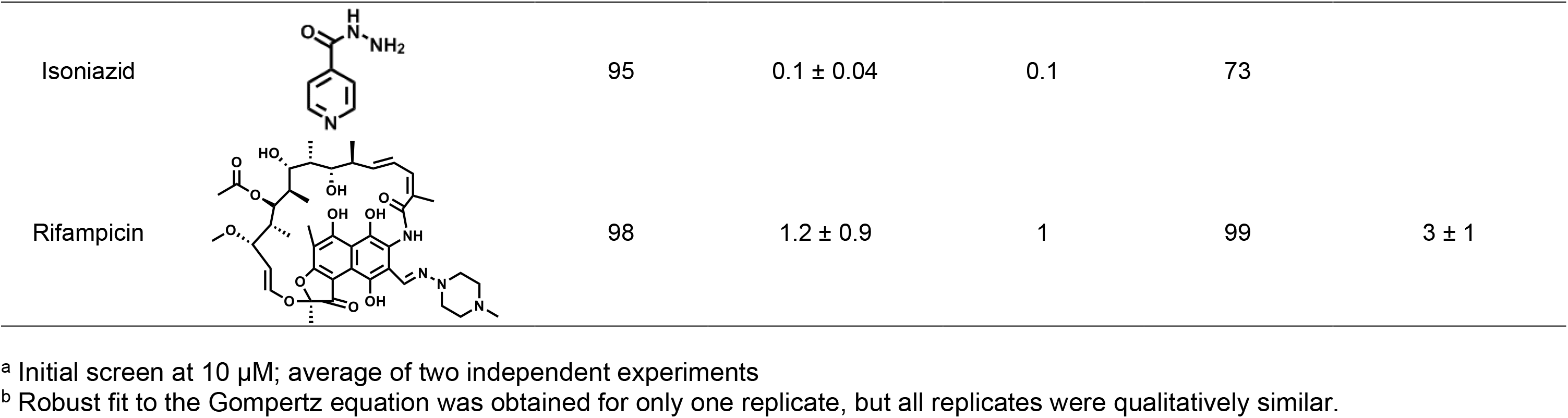
Activity of 1,2,3-triazole ureas against *Mycobacterium tuberculosis*

The morphological changes for cells treated with AA691, AA692, AA701, or AA702 were then processed and analyzed using the MorphEUS analysis pipeline and an existing reference drug set (Smith et al., 2020). The profile for each compound at a designated time point was individually applied onto the morphological space, constructed using 34 compounds with known molecular targets. Multiple classification trials (70) were performed for each analysis to determine the frequency of nearest neighbor connections. The resulting nearest neighbor frequency (connection strength) is highest among cells treated with drugs that target similar cellular components and pathways. This analysis therefore allows for classification of drug target(s) by determining the similarity in morphological response to drugs with known mechanisms of action.

## RESULTS

### Triazole urea compounds inhibit the growth of *Mtb* on glycerol and on cholesterol

To assess the activity of 1,2,3-triazole ureas against *Mtb* we screened a library of 192 compounds (Adibekian et al., 2011) at 10 μM for their ability to restrict growth. Screening on glycerol, the standard carbon source in laboratory growth medium, can lead to hit compounds that are inactive *in vivo* where glycerol is not a major carbon source (Pethe et al., 2010). We therefore tested for *Mtb* growth both on glycerol and on cholesterol, a nutrient source relevant *in vivo* (Wilburn et al., 2018). We defined hit compounds as those that decreased *Mtb* viability by ≥ 90% *vs*. vehicle-treated controls in both glycerol- and cholesterol-containing medium (**Figure 1, Table 1**). Based on this cutoff, the triazole ureas AA691, AA692, AA652, and AA321 were selected as hit compounds in our screen, for a hit rate of 2% (4 out of 192 compounds).

**Figure 1.**
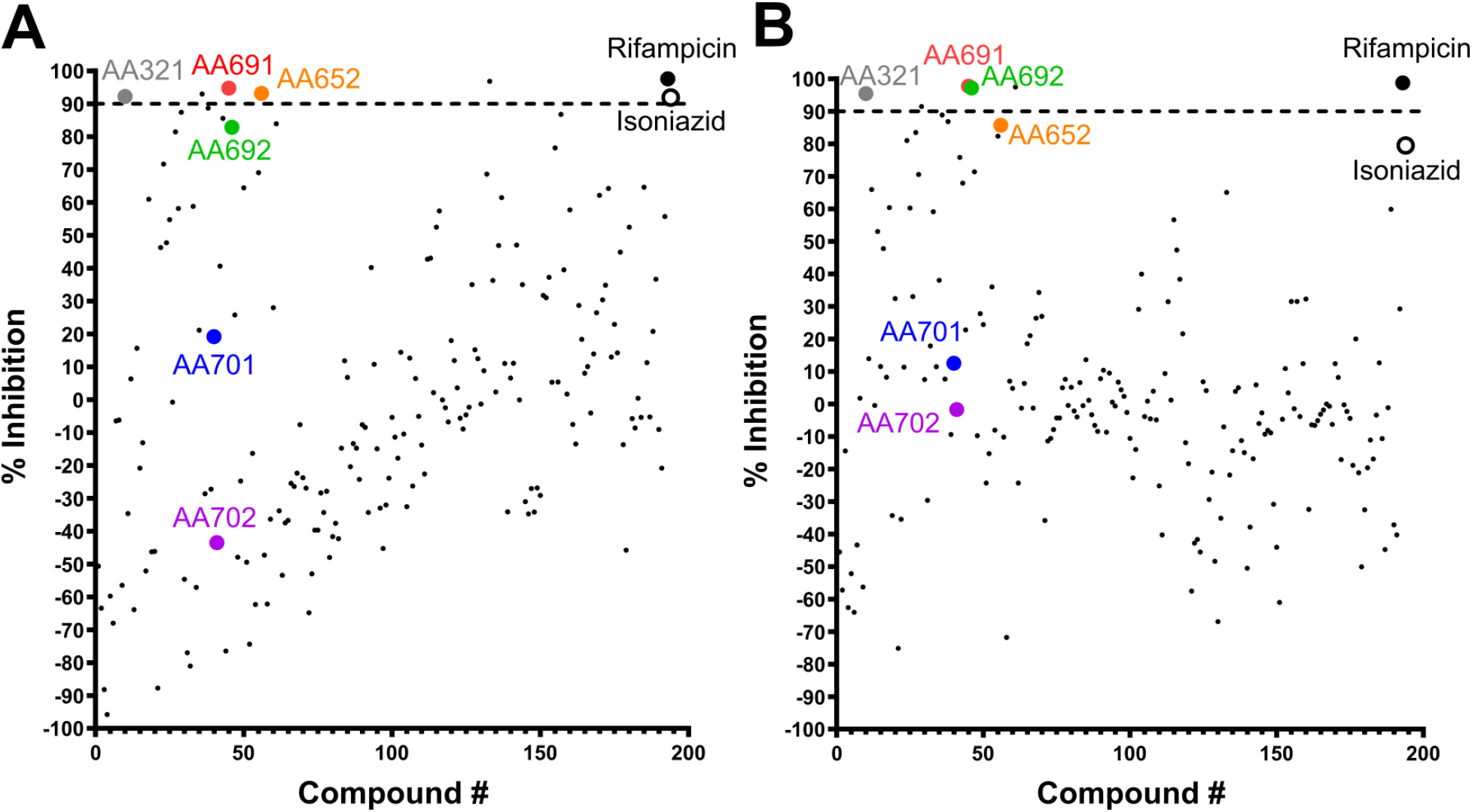
A triazole urea library yielded compounds that inhibit *Mtb* growth in both glycerol and cholesterol. Autoluminescent *Mtb* was subcultured to OD_600_ ~0.02 and incubated for 7 d with 10 μM compound in modified Roisin’s medium with A) glycerol or B) cholesterol as the sole carbon source. Percent inhibition was calculated by normalizing autoluminescence signal to DMSO vehicle-treated control. The Z′ scores were A) 0.59 and B) 0.49. Data shown are representative of 2 biological replicates.

Of the four hit compounds, AA691 and AA692 were of particular interest because of their relationships to two other compounds, AA701 and AA702. In this series AA692 serves as the parent structure (**Table 1**). AA691 is the largest and most hydrophobic compound and differs from AA692 in the substitution of a cyclohexanol for an isopropanol group at *C4* of the substituted triazole ring. AA701 and AA702 are successively smaller and differ from AA692 at the piperidine *C4*, with a methyl or no substitution, respectively. Interestingly, AA701 and AA702 exhibited successively lower activity than AA691 and AA692 in the initial screen (44% and −9% inhibition in glycerol; 32% and 20% in cholesterol, respectively.

We corroborated the initial screening results by measuring the minimum inhibitory concentrations (MIC) for AA691, AA692, AA701, and AA702 (**Figure S1**). The MICs were comparable by autoluminescence and visual inspection, confirming that these compounds affect *Mtb* viability and not just the function of the *luxABCDE* pathway. Consistent with the initial screen, AA691 and AA692 have low micromolar MICs, whereas AA701 and AA702 are ~4- and >10-fold less active (**Table 1**). In addition, we obtained the 1,4-regioisomers of AA691 and AA692 and found that these isomers were far less active (**Table 1**). For simplicity we hereafter use AA691 and AA692 to refer to the 2,4-regioisomers, except when making explicit comparisons to the 1,4-regioisomers.

### AA691 and AA692 restrict survival of replicating and non-replicating *Mtb*

During human infection, *Mtb* encounter diverse environmental stressors such as hypoxia and acidic pH (Gold and Nathan, 2017; Prosser et al., 2017). These factors can induce a state known as non-replicating persistence, in which *Mtb* remain metabolically active and grow and divide, but do not increase in number (Rittershaus et al., 2013). Previous studies have shown that *Mtb* serine hydrolases are active under hypoxia (Ortega et al., 2016; Tallman et al., 2016) and essential for intracellular pH homeostasis in acidic pH (Vandal et al., 2008; Zhao et al., 2015), suggesting that serine hydrolase inhibitors such as AA691 and AA692 could display antibacterial activity against both replicating and non-replicating *Mtb*. We cultured *Mtb* under hypoxia or in acidified medium (pH 5.0) to induce a state of non-replication upon simultaneous treatment with AA691 or AA692. We then measured autoluminescence and enumerated colony forming units (CFU) as a function of time and compound concentration. No *Mtb* autoluminesce was detected after incubation for 1 day under hypoxia, likely because the catalytic reaction that gives rise to luminescence requires oxygen. We instead determined cell viability under hypoxia only by enumerating CFU.

Due to the slow replication of *Mtb* in modified Roisin’s medium (doubling time ~4 days), we chose to determine compound activity over a total of 21 days and confirmed that all compounds tested were stable in growth medium over this period (**Figure S2**). We first confirmed that the selected culture conditions led to the predicted phenotypes for *Mtb*. Under conditions that support exponential growth, isoniazid was bactericidal after one doubling time, as expected (**Figure 2A**). Recovery of growth at later timepoints likely reflects acquisition of isoniazid resistance as previously documented in similar assays (Vilchèze and Jacobs, 2019). In contrast, under hypoxia the number of viable *Mtb* did not increase over time and isoniazid was not bactericidal at the same concentrations, confirming non-replication of *Mtb* and increased tolerance to isoniazid under these conditions. As a positive control, we confirmed that another front-line drug, rifampicin, remains bactericidal under hypoxia (**Figure 2B**).

**Figure 2.**
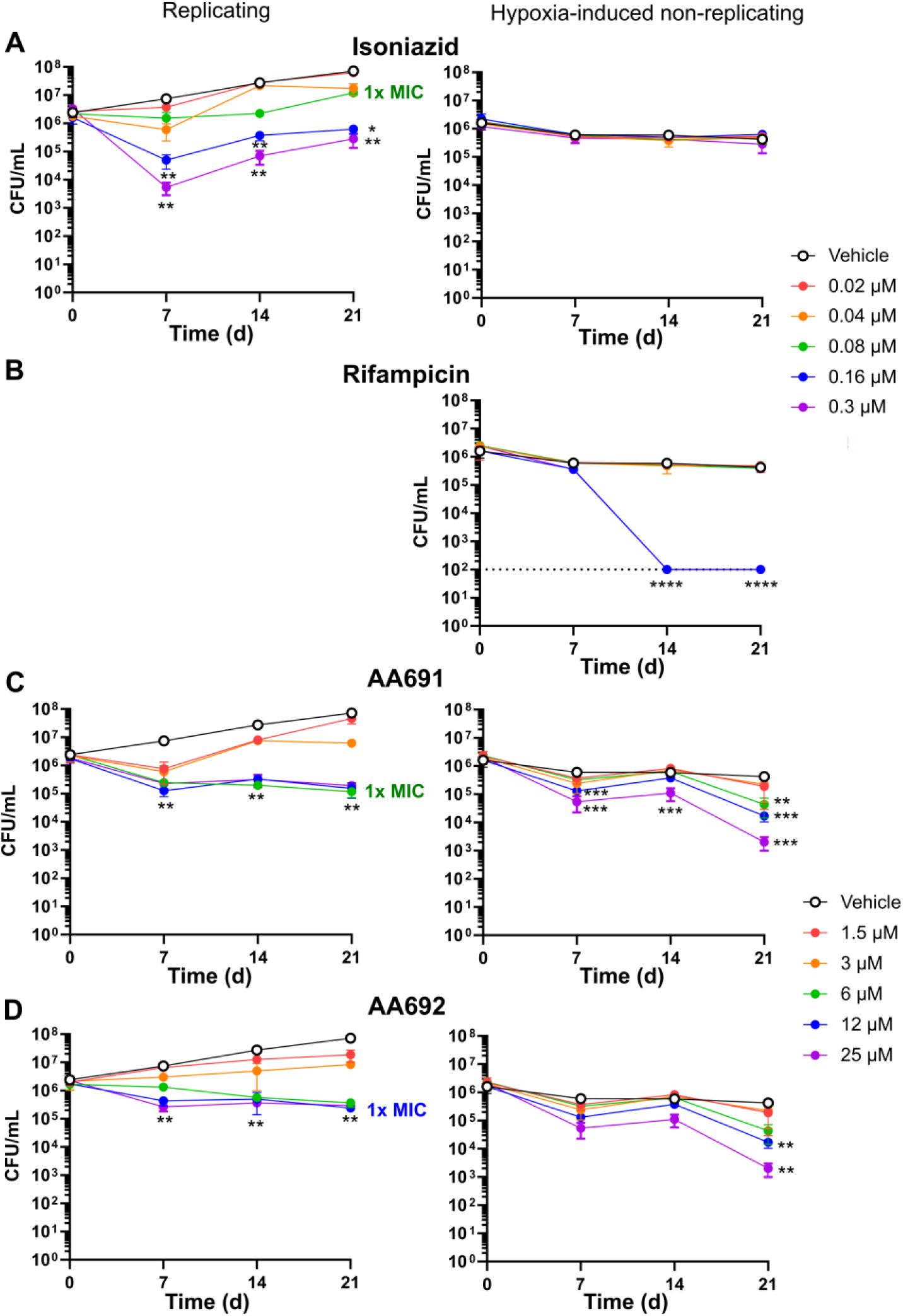
AA691 and AA692 are active against both replicating and hypoxia-induced non-replicating *Mtb*. *Mtb* was treated with A) isoniazid, B) rifampicin, C) AA691 or D) AA692 in modified Roisin’s medium containing glycerol and incubated under the indicated conditions. Bacteria were enumerated at each timepoint after 3-4 weeks incubation on 7H11 Middlebrook agar with10% OADC, 0.5% glycerol. * *p* < 0.05, ** *p* < 0.005, *** *p* < 0.0005, **** *p* < 0001 by one-way ANOVA with Dunnett correction for each timepoint vs. *t* = 0 (replicating) or vs. vehicle control (hypoxia-induced non-replicating). Data shown are the mean ± S.D. for 3 biological replicates.

In comparison AA691 and AA692 were weakly bactericidal against replicating *Mtb*. At their respective MICs, both AA691 and AA692 significantly decreased *Mtb* viability by 7 days and by up to ~1 log after 21 days **(Figure 2C, D)**. Higher concentrations of compound up to 4x MIC did not further decrease viability. When applied to *Mtb* just before inducing a non-replicative state by hypoxia, AA691 and AA692 had a more pronounced dose-dependent bactericidal effect with a 1-log decrease at 2X MIC and a 2-log decrease at 4X MIC compared to vehicle-treated controls after 21 days (**Figure 2C, D**). Similarly, AA691 and AA692 were also bactericidal when cultured at acidic pH, resulting in a significant decrease in *Mtb* autoluminescence by several-fold compared to vehicle-treated cells by day 11, while isoniazid activity was weaker but still significant (**Figure S3**).

### Rates of spontaneous resistance to hit compounds AA691 and AA692 are low

Genetic mutations that confer resistance to AA691 and AA692 could indicate encoded proteins that are involved in the mechanism of action. However, no colonies were visible after plating *Mtb* on 5X or 10X MIC of AA691 or AA692, suggesting that the rate of resistance is ~1×10^−9^ or lower. Although no resistant mutants were obtained, these experiments demonstrated that under the tested conditions the spontaneous rate of resistance to AA691 and AA692 compares favorably to that of isoniazid (**Figure S4**).

### ABPP with FP-biotin identifies a core active serine hydrolase proteome shared across multiple studies

We then pursued a biochemical approach to identify targets impacted by selected inhibitors. We have previously used competitive ABPP with Stable Isotope Labeling of Amino acids in Culture (ABPP-SILAC) to identify the serine hydrolase targets of triazole ureas in mammalian cells by quantitative mass spectrometry (Adibekian et al., 2011). Here, we first applied an analogous ABPP-SILAC approach to *Mtb* using the fluorophosphonate-biotin (FP-biotin) probe to detect the active serine hydrolase proteome (**Table S1**). We detected 105 proteins by ABPP-SILAC (**Table S2**) that were further refined to 56 proteins based on reproducibility of detection (**Table S3**). Interestingly, all proteins detected by ABPP-SILAC at pH 5.0 were also detected at pH 6.6, suggesting that the active serine hydrolase profiles are similar under both conditions.

Comparison of this set of active serine hydrolase proteome with those detected in two other ABPP studies in *Mtb* (**Figure 3**) (Ortega et al., 2016; Tallman et al., 2016) showed that among the 56 proteins, 44 were previously detected, supporting our cutoff for serine hydrolase annotation. Only two were not predicted as serine hydrolases based on Pfam annotation: the putative aldehyde dehydrogenase Rv0458 and the fatty acid-CoA ligase FadD2. The remaining 11 proteins we detected were mostly hypothetical conserved proteins of unknown function, but which had been bioinformatically annotated as serine hydrolases. Only one protein, the type I fatty acid synthase Fas, was detected by both the previous studies, but not by us. Similar to Ortega *et al.*, we found that the detected serine hydrolase proteome is enriched relative to the *Mtb* genome in the functional categories of lipid metabolism (16% vs. 6%) and intermediate metabolism and respiration (46% vs. 22%) (Ortega et al., 2016). In summary, the use of the fluorophosponate probe led to the identification of ~50 active serine hydrolases, 25 of which constitute a “core” proteome detected in all three studies. These results suggest that variations in ABP structure and experimental procedures result in distinct, but overlapping, inventories of active serine hydrolases in replicating *Mtb*.

**Figure 3.**
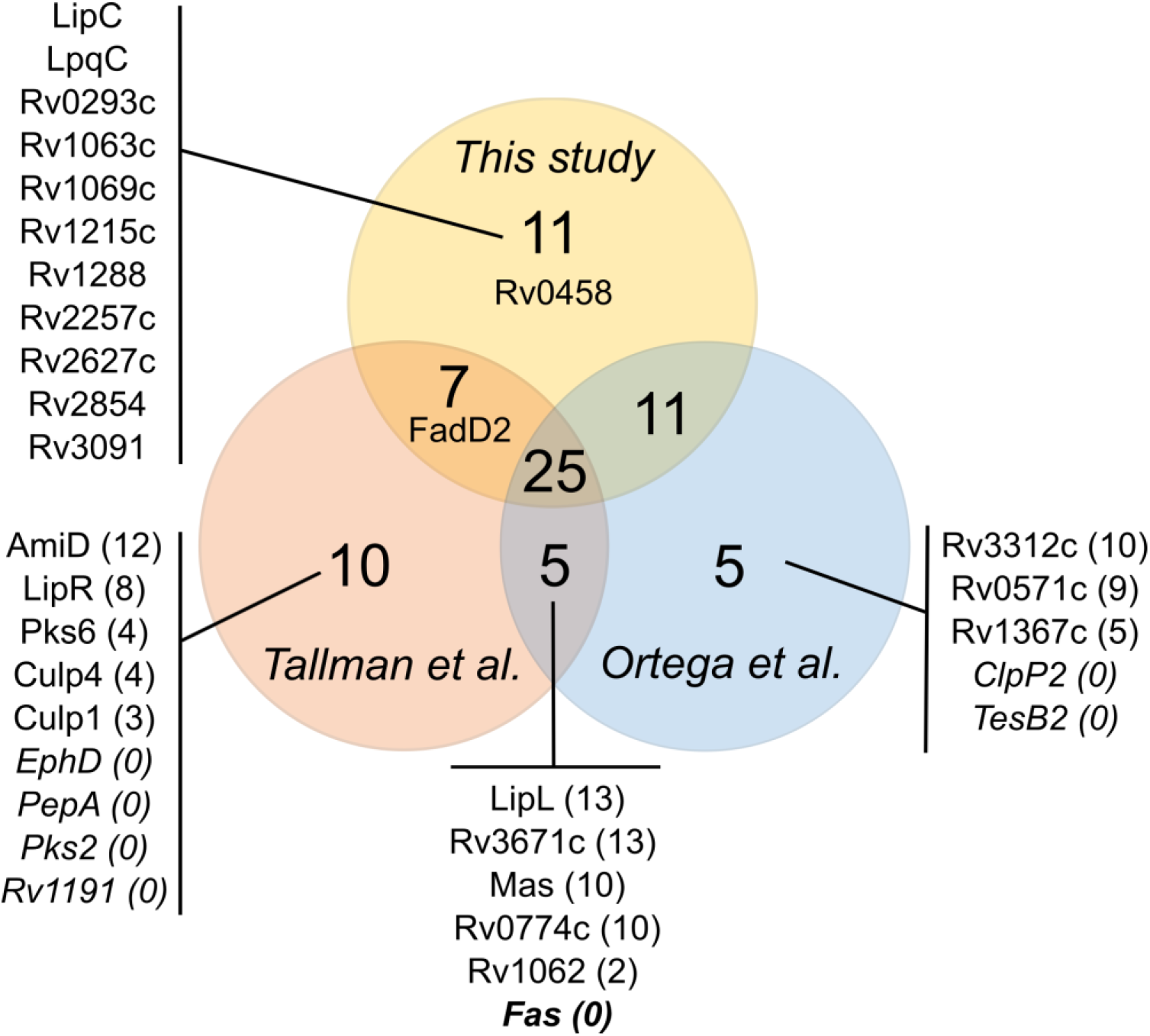
Activity-based protein profiling in *Mtb* by fluorophosphonate probes detects nearly 75 active serine hydrolase across multiple studies. The number in parentheses indicates the number of instrumental replicates in which a given serine hydrolase was detected in this study. Proteins without this annotation were detected in at least 14 of 15 instrumental replicates and thereby met our cutoff for annotation as a serine hydrolase. Proteins not detected in any instrumental replicates are in italics. The proteins Rv0458 and FadD2 met the detection cutoff, but are unlikely bona fide serine hydrolases and so were not counted.

### Preferential inhibition of individual serine hydrolases by AA692 over AA702 indicates high-priority targets

To quantify serine hydrolase inhibition by triazole ureas, we analyzed the competitive ABPP-SILAC experiments conducted at both pH 6.6 and pH 5.0, comparing compound- and vehicle-treated *Mtb* (**Figure S5**). We hypothesized that enzymes that are preferentially inhibited by AA691 and AA692 *vs*. the ~10-fold less active AA702 are more likely specific targets involved in the mechanism of action. Notably, AA691 was the more promiscuous inhibitor at pH 6.6 (**Table S3**), suggesting that the lower MIC of AA691 is due to the inhibition of additional targets, and that AA692, as the more specific inhibitor, would better illustrate the antibacterial activity of these compounds.

To delineate key targets likely involved in growth inhibition by AA692, we compared the difference in percent inhibition by AA692 *vs*. the inactive compound AA702 for each detected serine hydrolase (**Table S4**). This analysis yielded 11 (pH 6.6) and 15 (pH 5.0) prioritized targets, 8 of which overlapped (**Table 2**). This high degree of overlap suggests that AA692 retains not only antibacterial activity but also selectivity in targets under both conditions. Also, targets common to both conditions included most of those predicted as essential and detected as active under hypoxia. We therefore focused on the 11 prioritized targets identified at pH 6.6 as those most likely to be relevant to the antibacterial activity of AA692. Four have known or predicted functions in mycomembrane lipid biosynthesis: the mycolyltransferases FbpA and FbpB (also known as Ag85A and Ag85B) (Belisle et al., 1997); the thioesterase TesA (Alibaud et al., 2011; Chavadi et al., 2011); and the lipase Rv3802c (Parker et al., 2009). In addition to these lipid-related enzymes, Rv1730c is a predicted penicillin binding protein thought to function in maintaining peptidoglycan and thus in directly sustaining cell wall integrity.

**Table 2.** Prioritized serine hydrolase targets of AA692 identified by competitive activity-based protein profiling

| Protein name | Rv identifier | Prioritized targets |  | *In vitro* essential | Active under hypoxia | Identified as a target of** |  |  |  |
| --- | --- | --- | --- | --- | --- | --- | --- | --- | --- |
|  |  | pH 6.6 | pH 5.0 |  |  | EZ120 | lalistat | CyC<sub>17</sub> | THL |
| **AmiB2** | Rv1263 | • | • |  | a, b |  |  | • |  |
| **Rv1730c** | Rv1730c | • |  | 2 |  |  | • | • |  |
| **LipC** | Rv0220 | • | • |  |  |  |  |  |  |
| **Rv3591c** | Rv3591c | • | • |  | a |  |  |  |  |
| **FbpA** | Rv3804c | • | • | 2,4,5 | a | • |  | • |  |
| **Rv3802c** | Rv3802c | • |  | 1,2,3 | a |  |  |  |  |
| **FpbB** | Rv1886c | • | • |  | a | • |  |  |  |
| **Rv2627c** | Rv2627c | • | • |  |  |  |  |  |  |
| **TesA** | Rv2928 | • | • | 5 | a, b, c | • |  | • | • |
| **LipM** | Rv2284 | • | • |  | a, b, c | • | • | • | • |
| **Cut2** | Rv2301 | • |  |  |  |  |  |  |  |
| LipE | Rv3775 |  | • |  |  |  |  | • |  |
| LipG | Rv0646c |  | • |  |  |  |  |  |  |
| LipW | Rv0217c |  | • |  |  |  |  |  |  |
| Rv2854 | Rv2854 |  | • |  |  |  |  |  |  |
| LipN | Rv2970c |  | • |  | a, b |  | • | • |  |
| LipD | Rv1923 |  | • |  |  |  |  |  |  |
| BpoC | Rv0554 |  | • |  |  |  |  | • |  |

\*\* EZ120: Lehmann et al., 2017; lalistat: Lehmann et al., 2016; CyC<sub>17</sub>: Nguyen et al., 2017; THL: Ravindran et al., 2014
<sup>1</sup> Sassetti et al., 2003; <sup>2</sup> Griffin et al., 2011; <sup>3</sup> deJesus et al., 2017; <sup>4</sup> Bellerose et al., 2020; <sup>5</sup> FLUTE (https://orca2.tamu.edu/U19/)
<sup>a</sup> Ortega et al., 2016; <sup>b</sup> Tallman et al., 2016; <sup>c</sup> Ravindran et al., 2014

Among the other prioritized targets, Rv2627c is an uncharacterized protein; Rv3591c a possible hydrolase; LipC and LipM are esterases (Shen et al., 2012; Tallman et al., 2016) belonging to the hormone-sensitive lipase (HSL) family member proteins (*i.e*., Lip-HSL); AmiB2 is a probable amidase (a broad family that includes peptidoglycan-processing enzymes); and Cut2 (also known as Culp2) is a cutinase-like protein, one of seven Cut/Culp enzymes with *in vitro* esterase/phosopholipase activities (West et al., 2009). Together these findings suggest that AA692 inhibits several key serine enzymes involved in cell wall biosynthesis and thus enacts its antibacterial activity by disrupting the mycobacterial cell wall.

Of all the prioritized targets of AA692, the putative lipase Rv3802c is predicted essential in multiple studies in *Mtb* (DeJesus et al., 2017; Griffin et al., 2011; Sassetti et al., 2003) and confirmed essential in *M. smegmatis* (Meniche et al., 2009). Several other targets such as FbpA, TesA, and Rv1730c are predicted to be conditionally essential on restricted growth media (Bellerose et al., 2020; Griffin et al., 2011). In addition, FbpA, TesA, and Rv3802c are expressed and active under hypoxia (Ortega et al., 2016). The activity and essentiality of these targets under *in vitro* conditions similar to those used here further underscored their prioritization as targets of AA692.

### Biochemical assays and *in silico* molecular docking study validate ABPP-SILAC target identification and structure-activity relationships

To validate the ABPP-SILAC results, we next assessed inhibitor activity *in vitro* with several purified serine hydrolases. According to the ABPP-SILAC results, TesA and FbpA are preferentially inhibited by AA691 and AA692 compared to AA702, whereas Rv0183 is inhibited >95% by all three compounds. Purified TesA was preincubated with each compound at various inhibitor molar excesses (*x*_I_) and then subjected to either a substrate hydrolysis assay or a competitive ABP assay with the TAMRA-FP, a fluorescent ABP (Patricelli et al., 2001). A *x*_I50_ value of 0.5 is synonymous with a 1:1 stoichiometric ratio between the inhibitor and the lipolytic enzyme and is the highest level of inhibitory activity that can be achieved.

In both assays, AA691 and AA692 inhibited TesA in close stoichiometry (*x*_I50_ 0.8-1.8), in contrast with AA701 and AA702, which both exhibited >5-fold higher *x*_I50_ and thus lower relative inhibition (**Figure 4A, B**). We then used the competitive ABP assay to characterize the inhibition of FbpA. Similar to TesA, FbpA was significantly inhibited in a dose-dependent manner by AA691 and AA692, but not by AA701 or AA702 (**Figure 4A**). The regioisomers AA691-(1,4) and AA692-(1,4) showed intermediate potency against both enzymes. In contrast to these structure-activity relationships in the inhibition of TesA and FbpA, all tested compounds fully impaired Rv0183 activity (**Figure 4B**).

**Figure 4.**
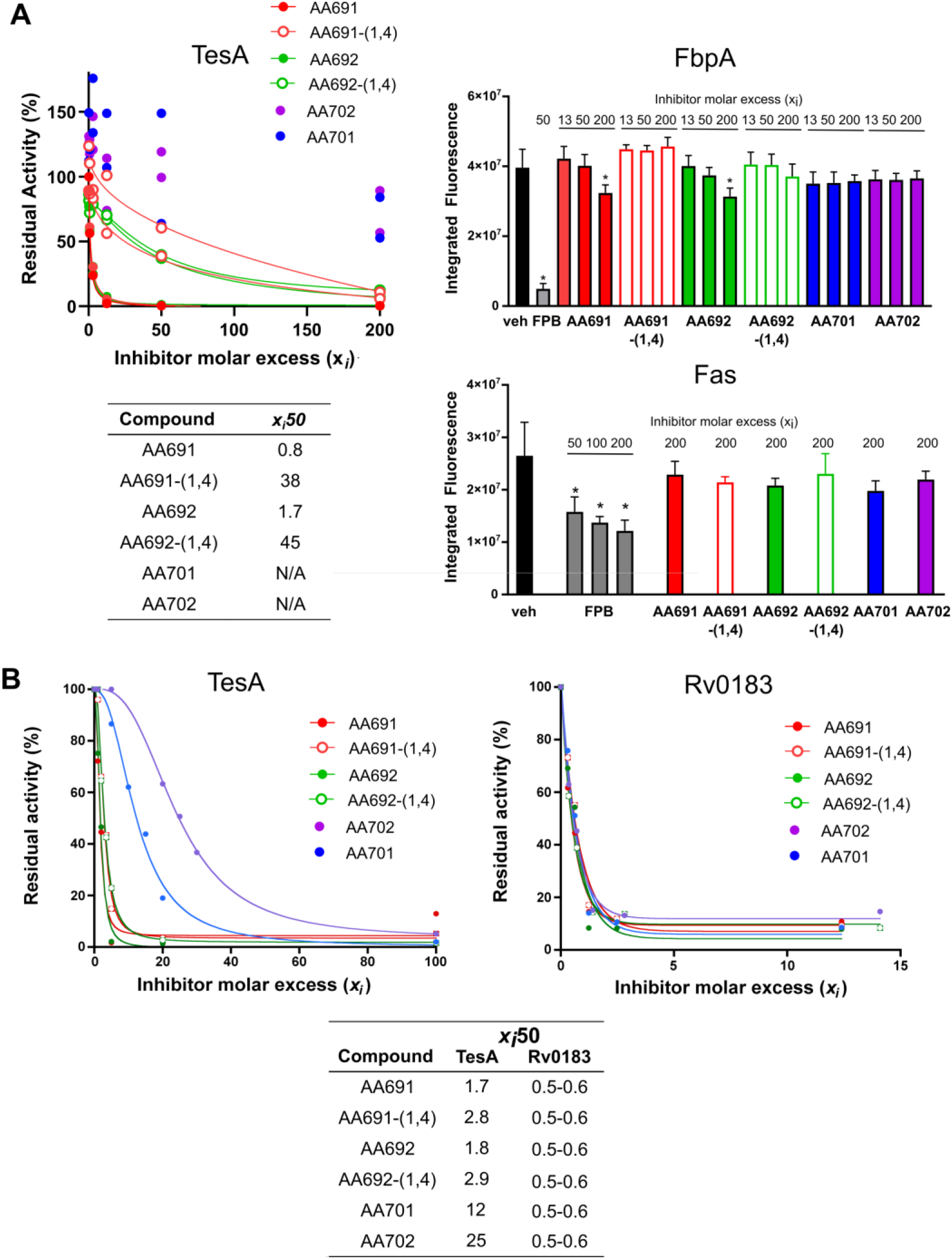
Inhibition of individual serine hydrolases recapitulates structure-activity relationships reported by competitive ABPP. Residual activity of purified serine hydrolases after incubation with the indicated compounds was determined by measuring A) FP-TMR labeling or B) residual activity via *p*NP-valerate hydrolysis followed colorimetrically for TesA or monoolein hydrolysis followed potentiometrically for Rv0183. The molar excess needed to reduce activity by 50% (*x*_I50_) was calculated by fitting the inhibition curve. Data shown are two independent experiments (A: TesA) or the average ± S.D. of 3 independent experiments (all others). * *p* < 0.05 by one-way ANOVA with Dunnett correction for a given concentration vs. vehicle-treated control.

We next sought to corroborate the observed biochemical activity of AA691, AA692, AA701 and AA702 using *in silico* docking to examine their predicted binding modes in the active sites of TesA and Rv0183. For TesA, docking provided possible explanations for the different inhibitory activities observed between the potent inhibitors AA691 and AA692 and the weak inhibitors AA701 and AA702 (**Figure 5A**). In these models AA691 and AA692 occupy the entire active site crevice of TesA and the carbonyl is at a favorable distance (2.1-2.2 Å) and orientation for forming a covalent bond with the catalytic Ser104 (**Figure 5A-B**). AA701 and AA702 adopt a similar binding mode when docked, but are farther from Ser104 (2.7 Å) (**Figure 5B**). AA691 obtained the most favorable binding interaction; AA702 achieved the least favorable (Δ*E* = −7.0, −6.7, −6.0, and −5.7 kcal/mol for AA691, AA692, AA701, and AA702, respectively). AA691 and AA692 are stabilized by an overlapping set of hydrophobic interactions (with <u>His36</u>, Ala37, <u>Met108</u>, Ser133, <u>Thr178</u>, Ile185, Ile210, His236 and Phe237; those unique to AA691 are underlined; **Figure 5D, E**). Binding of AA692 is also supplemented by two hydrogen bonds with Met105 and Cys132. Finally, the poses of AA701 and AA702 are almost superimposable, with a hydrogen bond to Ser133 for AA701 and similar hydrophobic interactions, but fewer contacts overall than for AA691 or AAA692 (**Figure 5F, G**).

**Figure 5.**
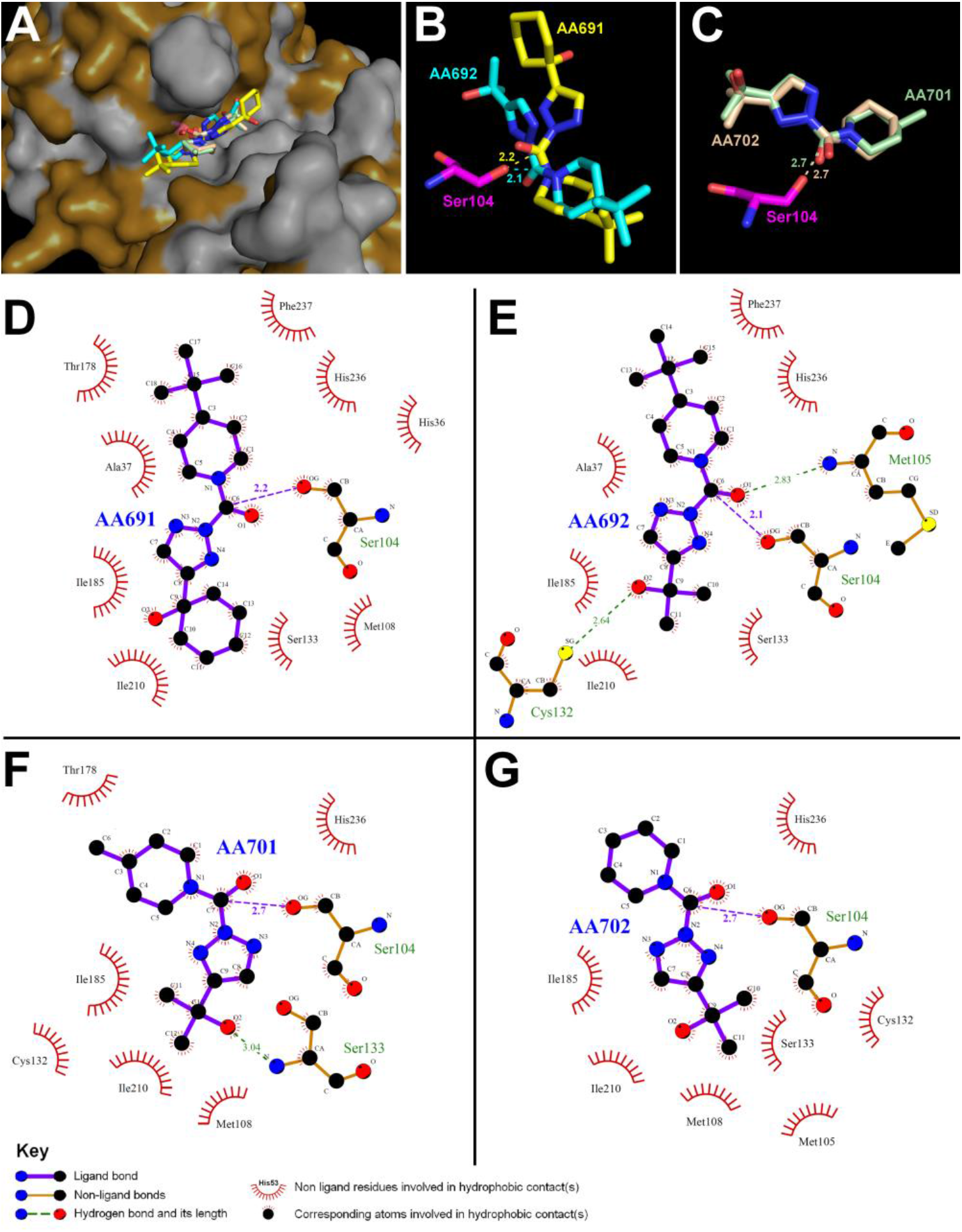
AA691 and AA692 make more contacts and are positioned closer to the catalytic serine than AA701 and AA702 in the TesA active site. A) *In silico* molecular docking of AA691, AA692, AA701 and AA702 into the crystallographic structure of TesA in a van der Waals surface representation. Hydrophobic residues are highlighted in *white*. Superimposition of the top-scoring docking position of B) AA691 (*yellow*) and AA692 (*cyan*) and C) AA701 (*pale green*) and AA702 (*wheat*) in the vicinity of the catalytic Ser104 (*magenta*). Structures were visualized with PyMOL using the PDB file 6FVJ. Ligplot+ analyses showing the ligand-protein interactions for D) AA691; E) AA692; F) AA701; and H) AA702 in the TesA active site with hydrogen bonds (*purple, green* dashed lines) and hydrophobic interactions (*red*) indicated.

In contrast, the computationally predicted binding modes for the compounds in Rv0183 were far more similar. All were predicted to adopt comparably productive orientations inside the enzyme active site (**Figure S6A-C**), with similar distances from the catalytic Ser110 (<2.5 Å) and similar predicted binding energy values (Δ*E* = −7.3 to −7.5 kcal/mol). Also, each inhibitor would be stabilized by largely the same hydrophobic interactions and hydrogen bonds (**Figure S6D-G**). Together, the predicted binding orientations, energies, and interactions in TesA and Rv0183 corroborate the biochemical inhibition data and the relative inhibitory potencies of the four inhibitors.

Given our hypothesis that AA692 is active against *Mtb via* the disruption of lipid and cell wall metabolism, we also purified Fas, as an essential fatty acid synthase that was the only serine hydrolase detected in previous studies (Ortega et al., 2016; Tallman et al., 2016), but not by us in any replicate. None of the compounds inhibited Fas significantly in the competitive ABP assay (**Figure 5A**). In addition, competition by biotin-FP was weak, showing that this probe does not efficiently label Fas and potentially explaining why Fas was not detected in our ABP profile.

### Morphological profiling confirms that AA691 and AA692 inhibit *Mtb* growth by disrupting cell wall synthesis

Although ABPP reveals proteins covalently modified by a given compound, it does not indicate which targets contribute to biological activity. To better understand the mechanism of action underlying AA692’s antibacterial activity, we next classified the responses of *Mtb* to these compounds using the recently developed morphological profiling platform for *Mtb* called MorphEUS (Smith et al., 2020). MorphEUS is based on the principle that drugs with similar mechanisms of action will induce similar changes in bacterial morphologies. Consistent with our target analysis, AA692 treatment caused morphological changes similar to those induced by cell wall synthesis inhibitors at both low and high dose (~0.5X and 5X MIC for AA692; **Figure 6A**). This pattern was corroborated by the results from AA691 at low dose. In contrast AA701 yielded a profile consistent with inhibition of protein synthesis with a weak connection to inhibition of cell wall synthesis that was more pronounced at high dose. The profiles resulting from AA702 most closely resembled protein synthesis inhibitors. Overall, MorphEUS analysis implicates cell wall synthesis most strongly in our hit compounds AA691 and AA692 and weakly or not at all in the less active compounds AA701 and AA702.

**Figure 6.**
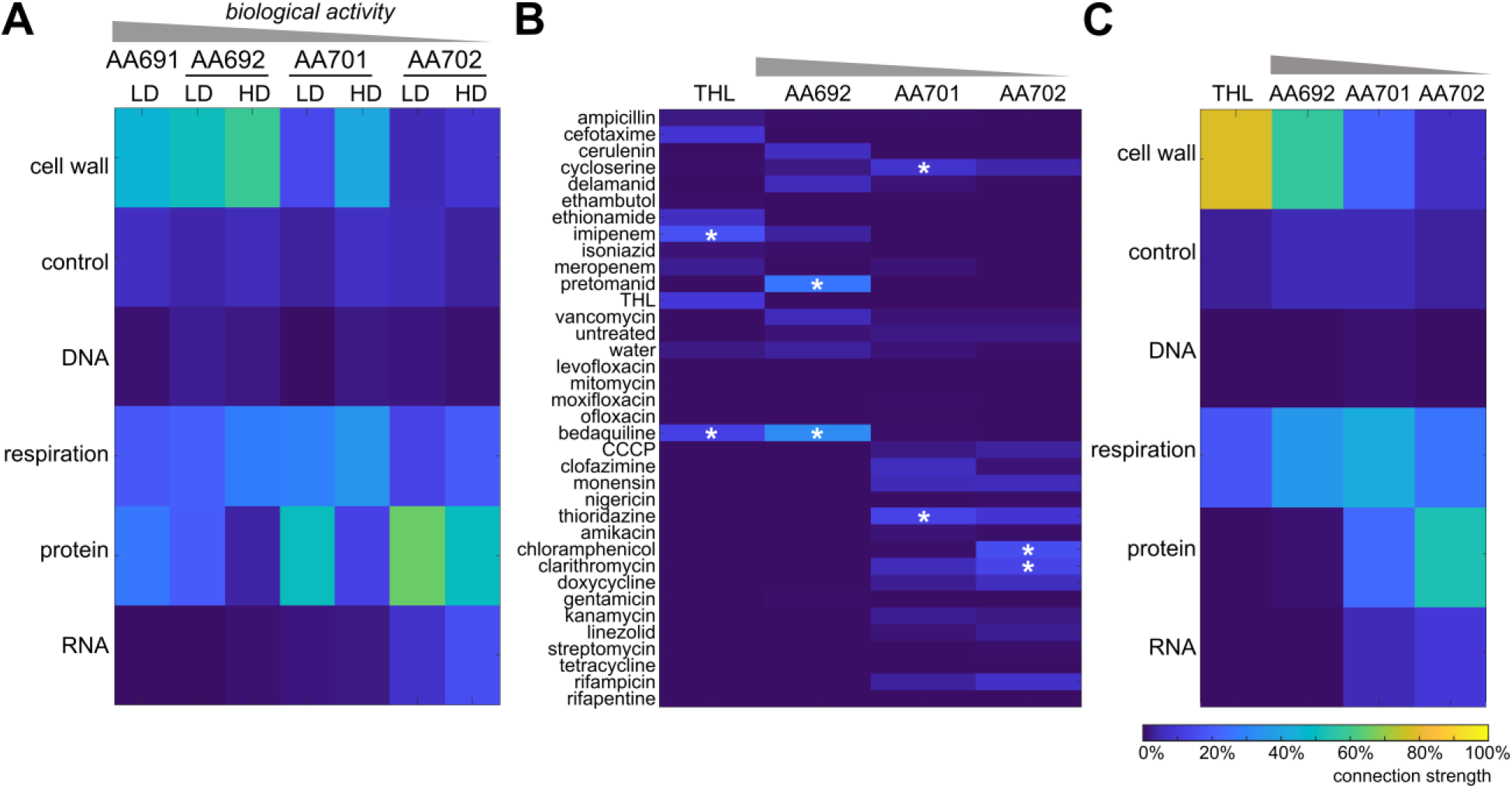
AA691 and AA692 cause morphological changes in *Mtb* similar to those induced by cell wall inhibitors. *Mtb* were incubated with 50 μM (low dose; LD) or 500 μM (high dose; HD) of the designated compounds and then stained for cellular membranes and the chromosomal nucleoid. Stained *Mtb* were imaged and analyzed for 25 morphological features. The profile for each compound at a given concentration was individually applied onto the morphological space constructed using 34 compounds with known mechanisms of action. The resulting nearest neighbor frequency (connection strength) based on A), C) broad categories corresponding to mode of action or B) individual compounds is highest among drugs that cause similar types of cellular change. A) shows data at both LD and HD; B) and C) are joint profiles from the simultaneous analysis of both LD and HD results (except for THL, for which the HD profile is shown). In B) the two most frequent neighbors for each compound are indicated with asterisks.

This conclusion is further supported by a review of individual compounds that represent the nearest neighbors of AA692 in the MorphEUS analysis: pretomanid and bedaquiline **(Figure 6B)**. Pretomanid acts in replicating Mtb by inhibiting the biosynthesis of essential mycolic acids. Although bedaquiline is an ATP-synthesis inhibitor, we and others have shown that the resulting downstream metabolic perturbation produces morphological changes that resemble those from cell-wall acting inhibitors (Mackenzie et al., 2020; Smith et al., 2020).

For comparison we also applied MorphEUS to tetrahydroplipstatin (THL; orlistat), a serine hydrolase inhibitor whose targets have been previously identified via ABPP (Ravindran et al., 2014). Two targets of THL, TesA and LipM, are also among the prioritized targets of AA692 (**Table 2**). THL shows a stronger connection to cell wall synthesis than AA692, likely because AA692 and related compounds are also connected to protein translation inhibitors, thereby reducing the apparent overall contribution of cell wall synthesis in the MorphEUS analysis (**Figure 6B, C**). Protein translation may be another pathway by which AA691 and AA692 exert their activity against *Mtb* since the translation inhibitor clarithromycin is among the nearest neighbors of both AA691 and AA692 at low dose (**Figure S7**). Alternatively, clarithromycin may have previously unrecognized effects on the cell wall that cause morphological effects similar to AA692. Overall, the results from MorphEUS analysis confirm disruption of cell wall biosynthesis as the direct consequence of AA691 and AA692 antibacterial activity, an assertion which correlates well with the lipid metabolism and cell wall synthesis enzymes identified by ABPP-SILAC as preferentially targeted by our hit compounds.

### AA691 and AA692 have mycobacteria-specific activity

Given the range of serine hydrolases targeted by all the 1,2,3-triazole ureas, we investigated whether our hit and related compounds might have activity against other bacteria. The antibacterial activity of AA691, AA692, AA701, and AA702 was assessed against *Escherichia coli*, *Staphylococcus saprophyticus*, and *Mycobacterium smegmatis* as representative Gram-negative, Gram-positive, and non-pathogenic mycobacterial organisms, respectively. AA691 and AA692 inhibited *M. smegmatis* growth with MICs of ~20 μM and ~30 μM respectively (**Figure 7A**). Consistent with their relative activities in *Mtb*, AA701 and AA702 had MICs >100 μM against *M. smegmatis.* In contrast, all four compounds had no detectable activity against *E. coli* or *S. saprophyticus* up to 100 μM (**Figure 7B, C**). These results support AA691 and AA692 as narrow-spectrum inhibitors.

**Figure 7.**
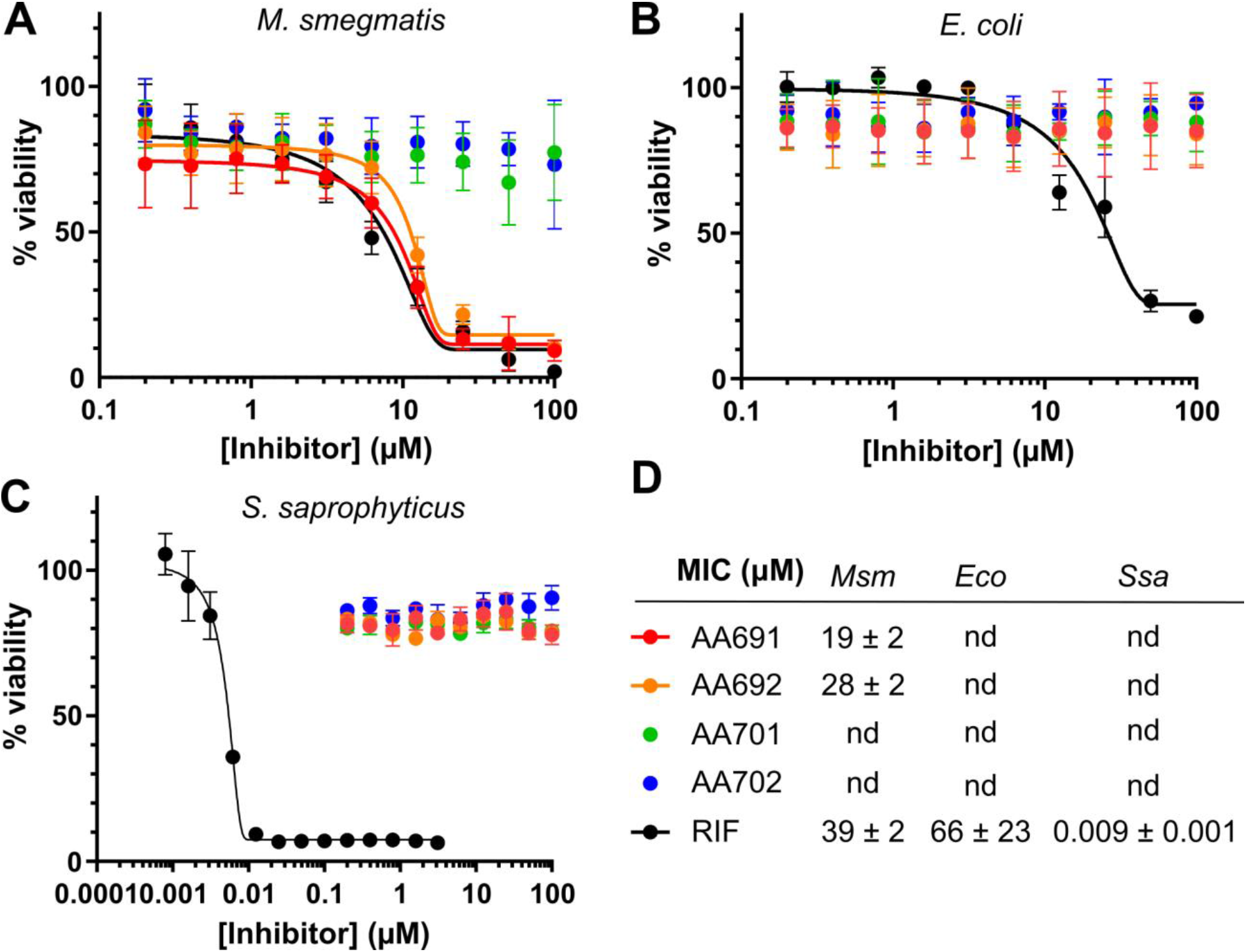
AA691 and AA692 do not have broad-spectrum antibiotic activity. A) Autoluminescent *M. smegmatis*, B) *E. coli* and C) *S. saprophyticus* were treated with the indicated compounds for ~3 doubling times for each respective bacterium. The viability of *was* determined by autoluminescence for *M. smegmatis* and by Bac-Titer Glo for *E. coli* and *S. saprophyticus*. Data shown are the mean ± S.D. of 3 technical replicates from one representative experiment in A)-C). MICs were determined by fitting the percent inhibition (versus DMSO vehicle-treated control) to the Gompertz equation and the mean ± S.D. from 3 independent experiments is reported in D); nd, not detected.

## DISCUSSION

In this study we evaluated a library of 1,2,3-triazole ureas for their ability to restrict the growth of *Mtb* in both glycerol- and cholesterol-containing medium. From this phenotypic screen we identified 4 hit compounds that inhibited *Mtb* growth ≥ 90% at 10 μM in both conditions for a hit rate of 2%. This hit rate is higher than that the <1% hit rate reported in other high-throughput anti-TB drug screens(Manjunatha and Smith, 2015). When tested against non-replicating *Mtb* induced by hypoxia or acidic pH, the hit compounds AA691 and AA692 both showed bactericidal activity, although the magnitude of this activity was lower under acidic pH compared to hypoxia. Most studies on serine hydrolase inhibitors have not reported activity against non-replicating *Mtb*. THL inhibits *Mtb* resuscitation upon return to normoxia (Kremer et al., 2005), but its activity against *Mtb* under hypoxia was not indicated. A study of β-lactams in combination with β-lactamase inhibitors found that none of the tested combinations killed *Mtb* under hypoxia (Solapure et al., 2013). Therefore, our compounds appear unique and our results provide foundational data on the potential of 1,2,3-triazole ureas to limit the survival of drug-tolerant non-replicating *Mtb*.

Spontaneous resistance mutations are commonly used to identify and confirm target proteins and pathways that underlie inhibitor activity, However, resistant mutants to the hit compounds AA691 or AA692 were not obtained here, suggesting that the multi-target nature of these inhibitors prevent the bacteria from developing any heritable resistance mechanism. Accordingly, competitive ABPP enables identification of proteins preferentially targeted by the hit compound AA692 vs. the antibacterially inactive compound AA702. Inhibition assays and *in silico* docking with individual targets validated our competitive ABPP results, showing that TesA and FbpA are preferentially inhibited by AA691 and AA692 while Rv0183 is efficiently inhibited by all compounds.

In contrast with our study, the targets of other serine hydrolase inhibitors such as THL, lalistat, EZ120, and CyC_17_ were identified without comparison to the targets of a related inactive compound (Lehmann et al., 2016, 2018; Nguyen et al., 2017; Ravindran et al., 2014). AA691 and AA692 share multiple targets with these serine hydrolase inhibitors, including the conditional essential enzymes FbpA, TesA, and Rv1730c (**Table 2**). Given this overlap, does our comparative analysis produce distinct conclusions? To address this question, we performed an additional analysis based on the activity of AA692 alone, rather than in comparison to AA702. In this alternate approach, the same number of prioritized targets (11) resulted from applying a cutoff of >60% inhibition by AA692. Five overlapped (TesA, Rv1730c, LipC, LipM, AmiB2) with prioritized targets from our comparative analysis (**Table 2**), but the other six serine hydrolases were excluded due to a high degree of inhibition by AA702 (Rv0183, LipG, LipH, LipO, AmiC, Rv0293c). Four of these excluded targets— Rv0183, the phospholipase/thioesterase LipG (Santucci et al., 2018), the esterase LipH (Canaan et al., 2004), and the putative lipase LipO)— were also identified as direct targets of the alkyne-modified derivatives of lalistat, THL, or EZ120. Our comparison of AA692 and AA702 suggests that these enzymes are non-selectively targeted by active and inactive antibacterial compounds and that their inhibition may thus be unrelated to restricting *Mtb* growth or survival.

While our comparative analysis provides a short list of key targets, ABPP reports on an inhibitor’s biochemical effects and not on which targets are most important to an inhibitor’s biological activity. For the six enzymes prioritized exclusively by comparing AA692 and AA702 target profiles, their contributions to the activity of AA692 ultimately depends on the vulnerability of each target, *i.e.*, the degree to which they must be inhibited to affect bacterial growth or survival. This has been addressed in mycobacteria on a limited scale using conditional proteolysis (Wei et al., 2011), but to our knowledge no data are yet publicly available for any of our prioritized targets. Moreover, target vulnerability may change when other targets are simultaneously inhibited; these probable effects have yet to be explored experimentally.

To obtain further clues about the antibacterial mechanism of action we pursued morphological profiling as a newly developed, independent approach that classifies compounds by comparison to well characterized inhibitors. By tracking morphological changes that occur in *Mtb* after compound treatment, we were able to classify our inhibitors according to similar morphological shifts. The MorphEUS analysis revealed that compounds with weak antibacterial activity closely resemble protein synthesis inhibitors while AA691 and AA692 resemble cell wall synthesis inhibitors. Such comparisons between active and inactive antibacterial compounds are integral to narrow down the mechanism of action, especially with multi-target inhibitors that do not easily yield resistant mutants and are thus refractory to traditional analysis of resistance mutations. Overall, these results strongly support that AA691 and AA692 disrupt cell wall synthesis by inhibiting lipid metabolism and cell wall synthesis enzymes identified by ABPP and confirm that ABPP identifies protein targets that are relevant to a compound’s mechanism of action.

Despite the broad targeting of serine hydrolases by AA692 and related triazole urea compounds, AA692 exhibited appreciable antibacterial activity only against mycobacteria compared to other gram positive and negative bacteria. This selectivity might be related to the unique composition of the mycobacterial cell envelope (Brennan and Nikaido, 1995) and the associated ability of these small hydrophobic compounds to cross the lipid-rich cell wall. This hypothesis is relevant given that serine hydrolases preferentially targeted by AA692 have lipid- and cell wall-related functions. However, given that 1,2,3-triazole ureas have been used successfully to inhibit serine hydrolases in live *S. aureus* (Chen et al., 2019), we favor the hypothesis that the hundreds of predicted mycobacteria serine hydrolases, many of which are annotated as essential, compared to the dozens so far detected in *E. coli* or *Staphylococcus* species (Keller et al., 2020; Shamshurin et al., 2014), make mycobacteria more susceptible to serine hydrolase inhibitors.

Based on both ABPP and microbiological analyses, all of the serine hydrolase inhibitors characterized in mycobacteria to date likely act by inhibiting cell wall synthesis via the inhibition of multiple enzymes. This is consistent with the large number of active serine hydrolases in *Mtb* as noted above and their enrichment in the functional categories of lipid metabolism and intermediate metabolism and respiration. The lack of resistance to AA691 and AA692 that we observed in our TesA overexpression strain is thus not surprising since overexpression of a single target enzyme is unlikely to generate a high level of resistance to a multi-target inhibitor.

The utility of triazole urea compounds as preclinical candidates against *Mtb* is speculative. The low selectivity index for AA691 and AA692 (**Figure S8**) advises against their use as preclinical candidates or in infection models of TB. However, triazole ureas that are specific for individual serine hydrolases have been developed for mammalian cells (Adibekian et al., 2011; Hsu et al., 2013b, 2013a), suggesting that the same is possible for *Mtb*. One advantage of triazole ureas is that their synthesis is straightforward, requiring only three steps to the finished product. This synthetic simplicity could facilitate library expansion into further structure-activity relationships to improve activity and selectivity while lowering toxicity. More generally, our observations support the serine hydrolase family as a promising set of targets and serine hydrolase-targeted inhibitor libraries as well positioned to exploit these pharmacologically vulnerable enzymes. Notably, in addition to triazole ureas, carbamates and aza-lactams have also proved versatile scaffolds for discovering and optimizing chemical probes and inhibitors for serine hydrolases. These libraries and others provide additional opportunities to exploit the screening and characterization platform that we have described here.

The results from our study confirm a limitation of the competitive ABPP approach: The detection of targets relies on (assumed) promiscuity of the competing ABP label. This contrasts with the majority of ABPP studies that have used alkyne derivatives of inhibitors to identify targets directly. However, not all inhibitors are amenable to chemical modification to add a handle such as an alkyne and although the addition of an alkyne is expected to be minimally perturbative, the biological activity of modified inhibitors must be confirmed. Thus, the competitive ABPP approach offers ease and versatility since hit compounds do not need to be adapted and re-validated. Competitive ABPP does require attentive matching of inhibitor libraries and probe to ensure that targets will be accurately captured by the probe. While matched libraries have been developed most extensively for serine hydrolases and applied using the fluorophosphonate probe, new ABPs are continually being developed to target additional chemistries on proteins, widening the scope for competitive ABPP in inhibitor discovery and characterization.

## AUTHOR CONTRIBUTIONS

M.L., H.V.P., A.B.C., S.C., B.B.A., B.F.C., and J.C.S conceived the study and designed the experiments; H.V.P. performed the compound screen; M.L. performed the microbiological assays; H.V.P., M.L. and A.B.C. performed the activity-based probe profiling; M.L. and I.M. performed the enzyme purification and inhibition assays; J.-F.C. performed the computational docking; T.C.S. performed the morphological profiling. M.L., A.B.C., I.M., S.C., T.C.C., J.C.S. analyzed the data. M.L., H.V.P., A.B.C., T.C.S., I.M., J.-F.C., S.C., B.B.A., and J.C.S. wrote and edited the manuscript.

## Notes

### Competing Interest Statement

The authors have declared no competing interest.

