## Supplemental Figures and Methods for "A competitive activity-based protein profiling platform yields cell wall synthesis inhibitors active against replicating and non-replicating *Mycobacterium tuberculosis*"

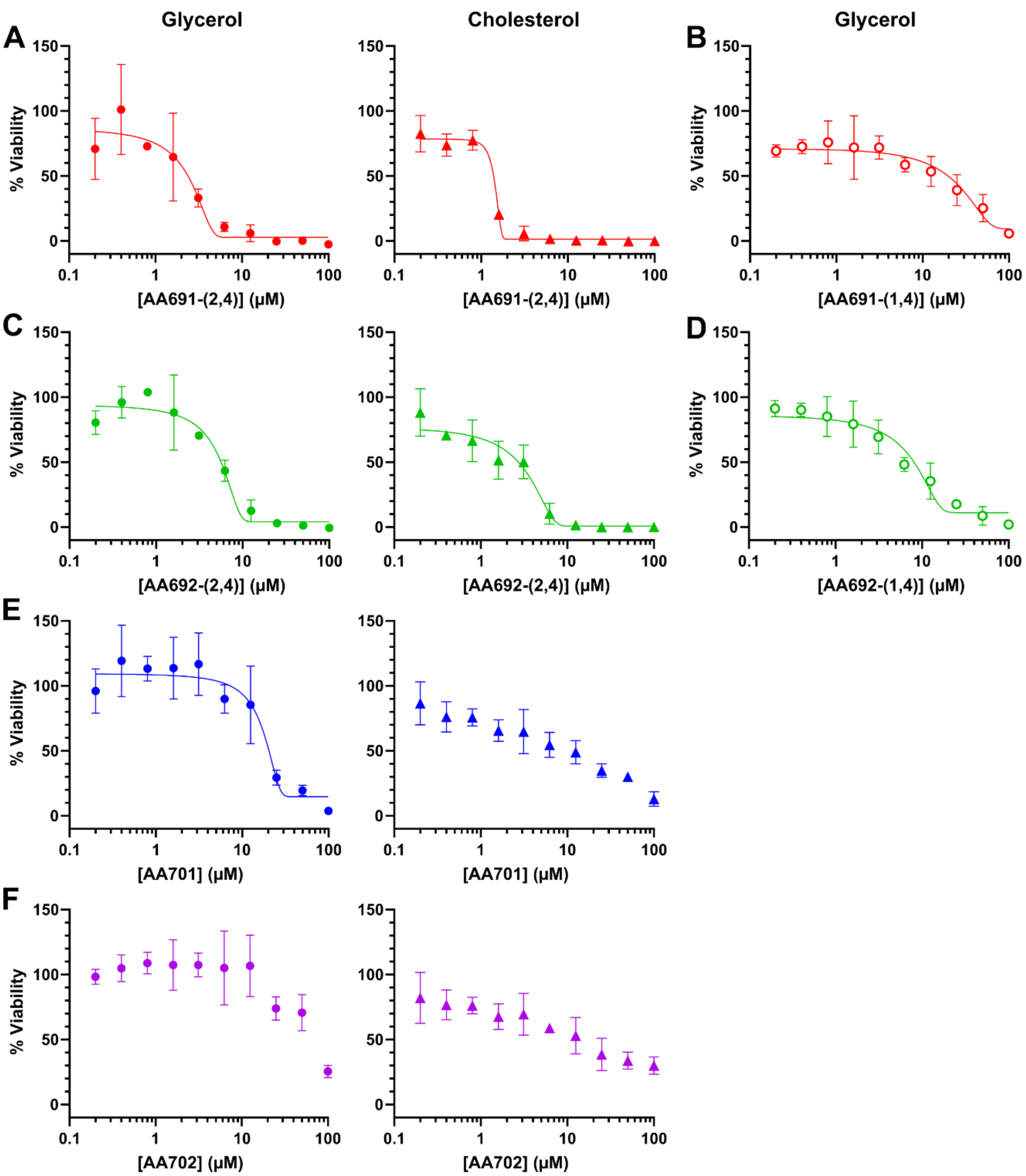


**Figure S1. Minimum inhibitory concentrations confirm structure-activity relationships.** Autoluminescent *Mtb* was treated with A) AA691, B) AA691-(1,4), C) AA692, D) AA692-(1,4), E) AA701, or F) AA702 in modified Roisin's medium containing glycerol or cholesterol as indicated. Each data point is the mean ± S.D. of 3 technical replicates from a single experiment; data shown are representative of three independent experiments. MICs were determined by fitting the percent viability (versus DMSO vehicle-treated control) to the Gompertz equation.

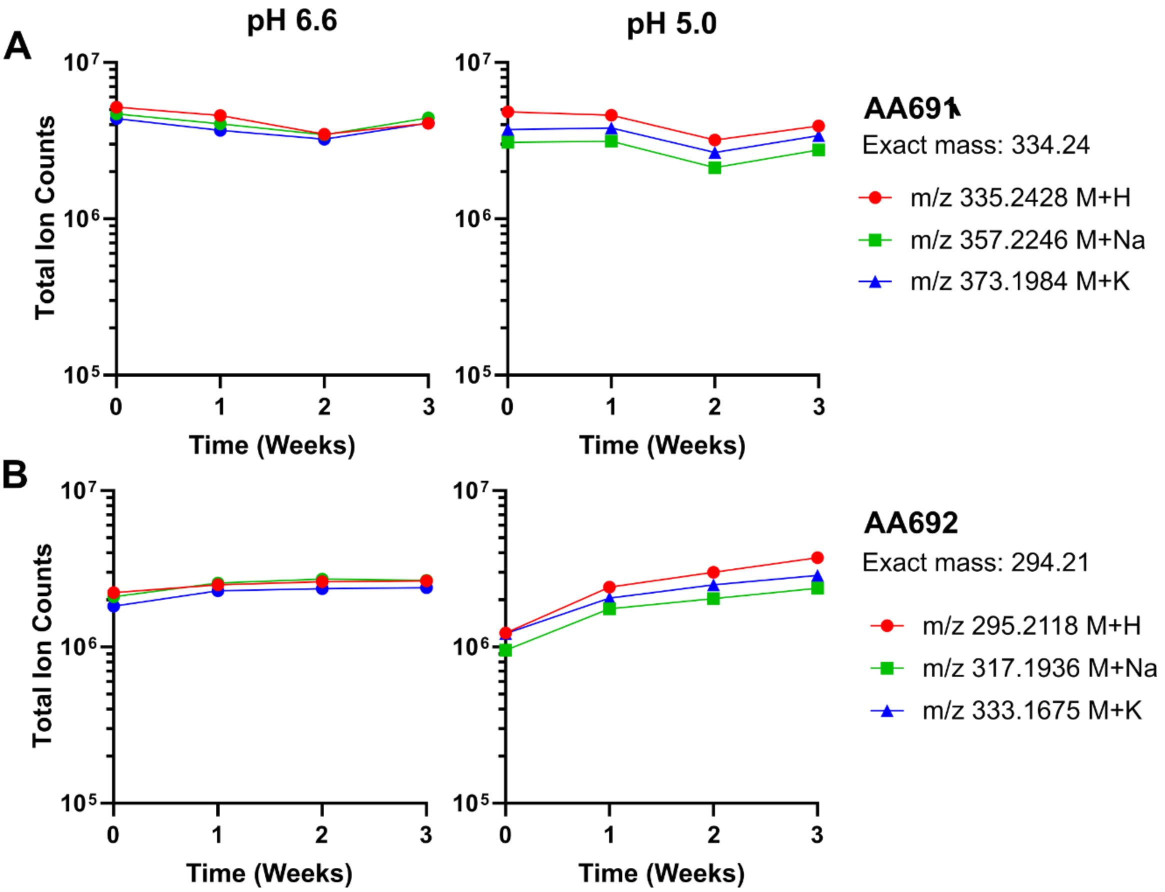


**Figure S2. AA691 and AA692 are stable in modified Roisin's medium.** A) AA691 or B) AA692 at a final concentration of 10 µM was incubated in modified Roisin's medium containing glycerol at pH 5.0 or pH 6.6 for 0, 1, 2, or 3 weeks at 37 °C. Samples were analyzed by liquid chromatography mass spectrometry. The *m/z* peaks corresponding to $[M + H]^+$, $[M + Na]^+$, and $[M + K]^+$ were manually assigned. Data are from individual samples at each time point from a single experiment.

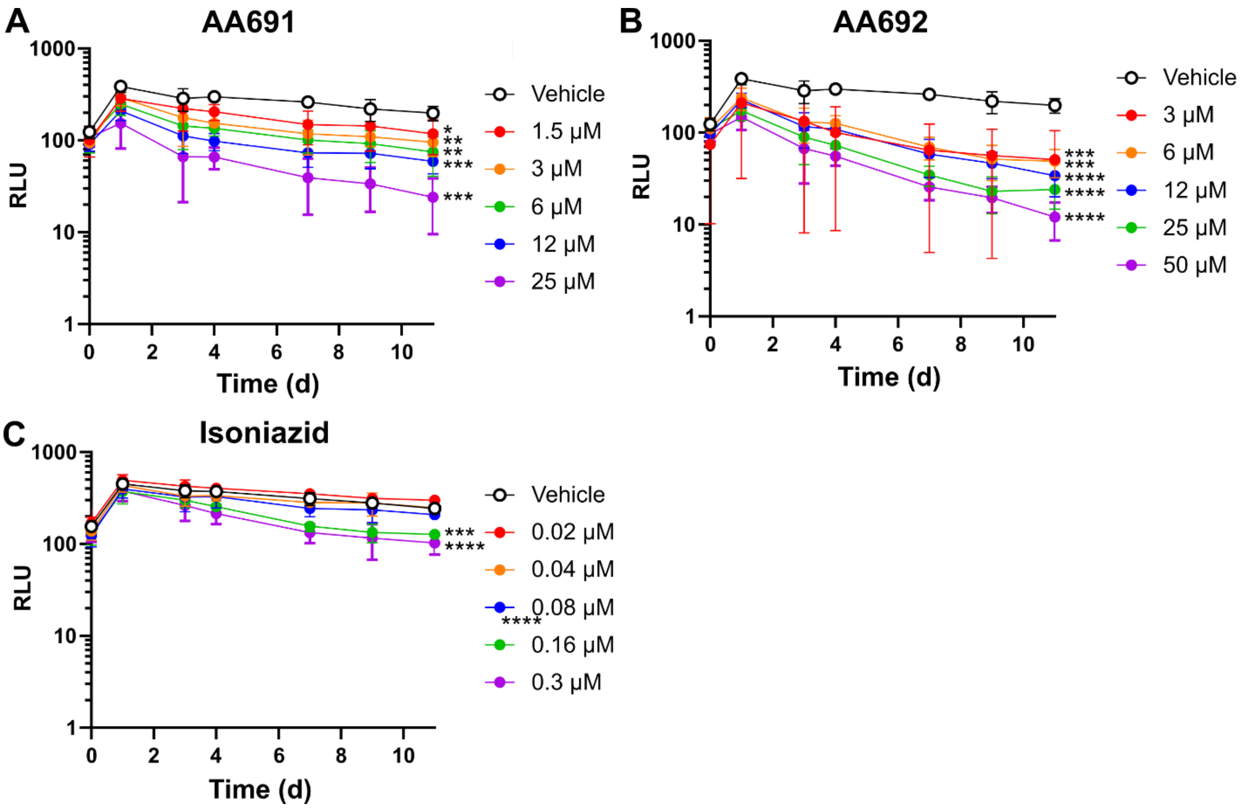


**Figure S3. AA691 and AA692 are active against acidic pH-induced non-replicating *Mtb*.** Autoluminescent *Mtb* were incubated in modified Roisin's medium at pH 5.0 for 3 days as an adaptation step before treatment with A) AA691, B) AA692, or C) isoniazid. For assays in acidic pH, *Mtb* was subcultured to a higher $OD_{600}$ of 0.1 (vs. 0.02 at pH 6.6) due to limits of luminescence when bactericidal activity was observed. * $p < 0.05$, ** $p < 0.005$, *** $p < 0.0005$, **** $p < 0001$ by one-way ANOVA with Dunnett correction for each timepoint vs. vehicle-treated control on day 11. Comparisons at other timepoints were not significant. Data shown are the mean ± S.D. of 3 biological replicates.

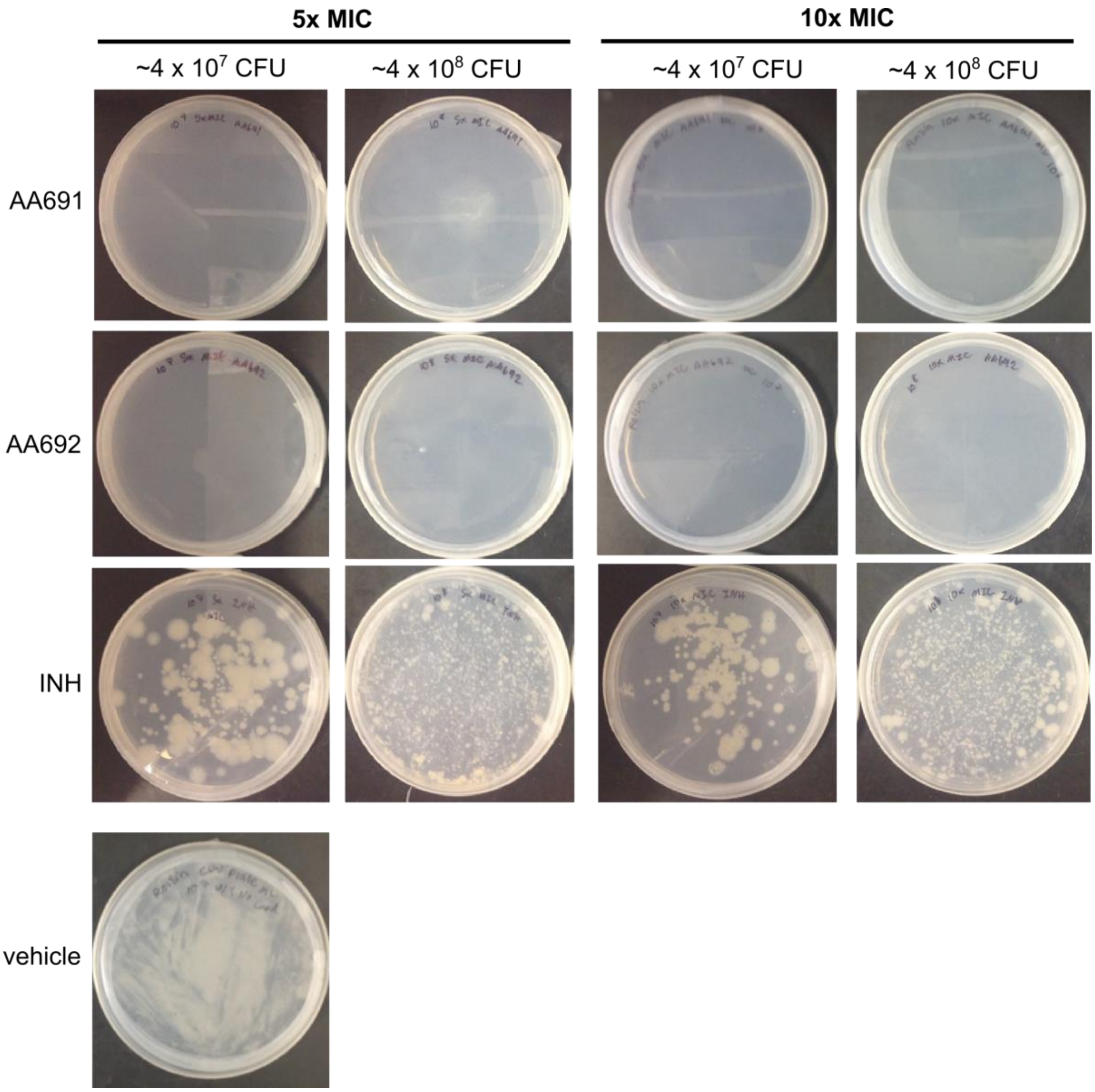


**Figure S4. The spontaneous resistance rate against AA691 and AA692 was lower than 1x10$^{-8}$.** *Mtb* at the indicated estimated CFU were plated on Roisin's modified medium agar containing glycerol and 5X or 10X MIC AA691, AA692, or INH (where 10X MIC corresponded to 60 μM AA691, 120 μM AA692, or 0.8 μM INH). Data shown are representative of two independent experiments.

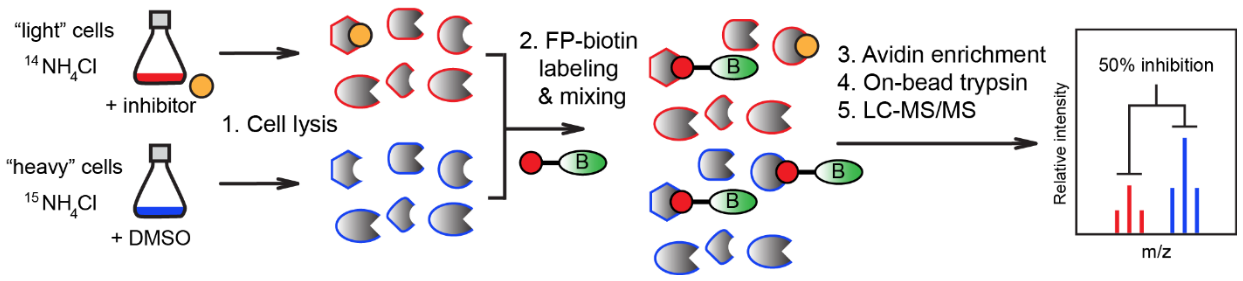


**Figure S5. ABPP-SILAC enables quantitative determination of serine hydrolase inhibition following treatment of whole cells with compounds.** *Mtb* were cultured in "light" ($^{14}N$) or "heavy" ($^{15}N$) medium and treated with inhibitor and vehicle control, respectively. Following cell lysis, labeling with FP-biotin, the samples were mixed, enriched, and analyzed by LS-MS/MS. Percent inhibition was determined by comparing the "light" vs. "heavy" peptides (compound treated vs. untreated) assigned to each protein detected.

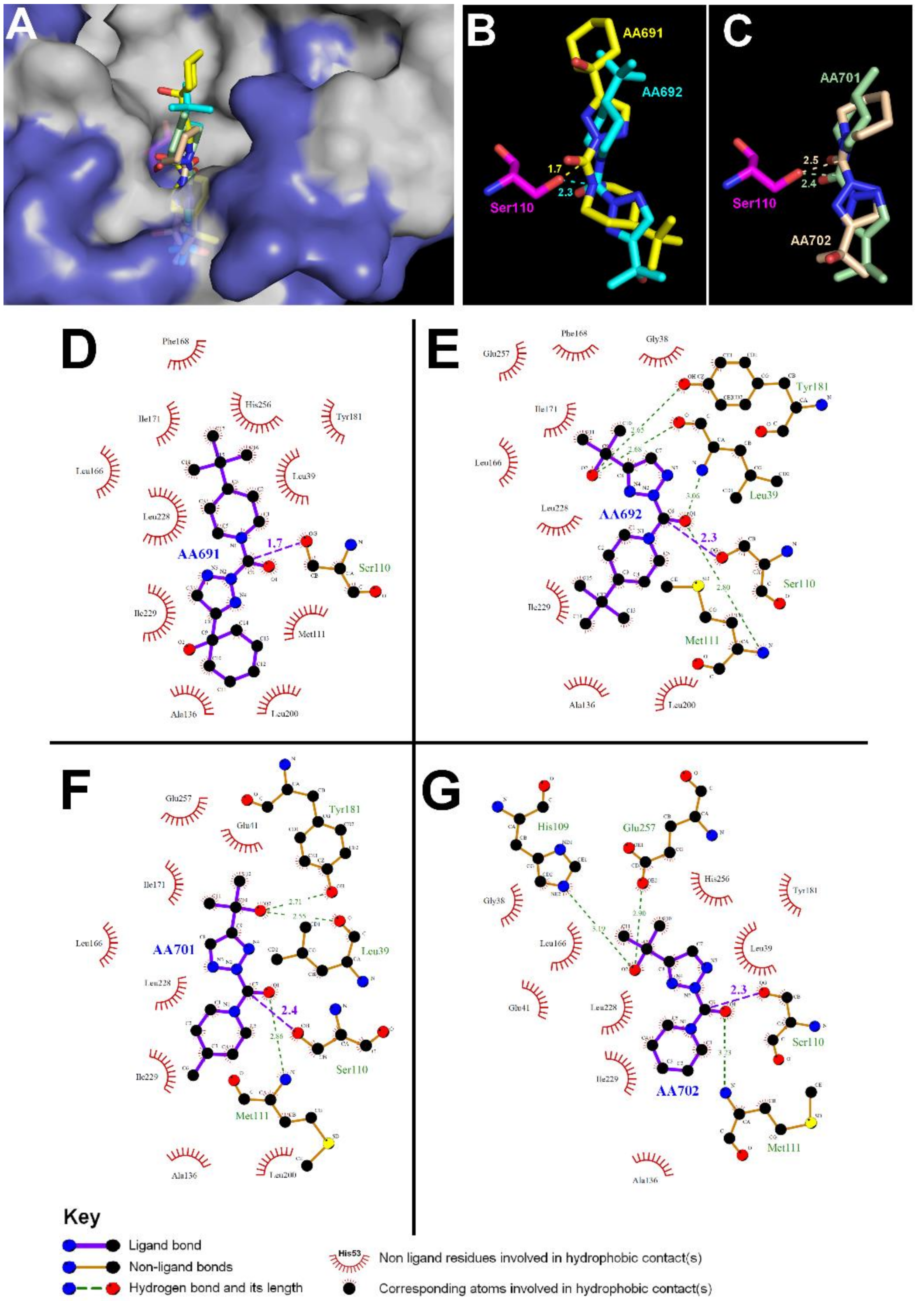


**Figure S6**. **AA691, AA692, AA701, and AA702 adopt similar poses in the Rv0183 active site.** A) *In silico* molecular docking of AA691, AA692, AA701 and AA702 into the crystallographic structure of Rv0183 in a van der Waals surface representation. Hydrophobic residues are highlighted in *white*. Superimposition of the top-scoring docking position of B) AA691 (*yellow*) and AA692 (*cyan*) and C) AA701 (*palegreen*) and AA702 (*wheat*) in the vicinity of the catalytic Ser110 (*magenta*). Structures were drawn with PyMOL using the PDB file 6EIC. Ligplot$^{+}$ analyses showing ligand-protein interactions for D) AA691; E) AA692; F) AA701 and H) AA702 in the Rv0183 active site with hydrogen bonds (*purple, green* dashed lines) and hydrophobic interactions (*red*) indicated.

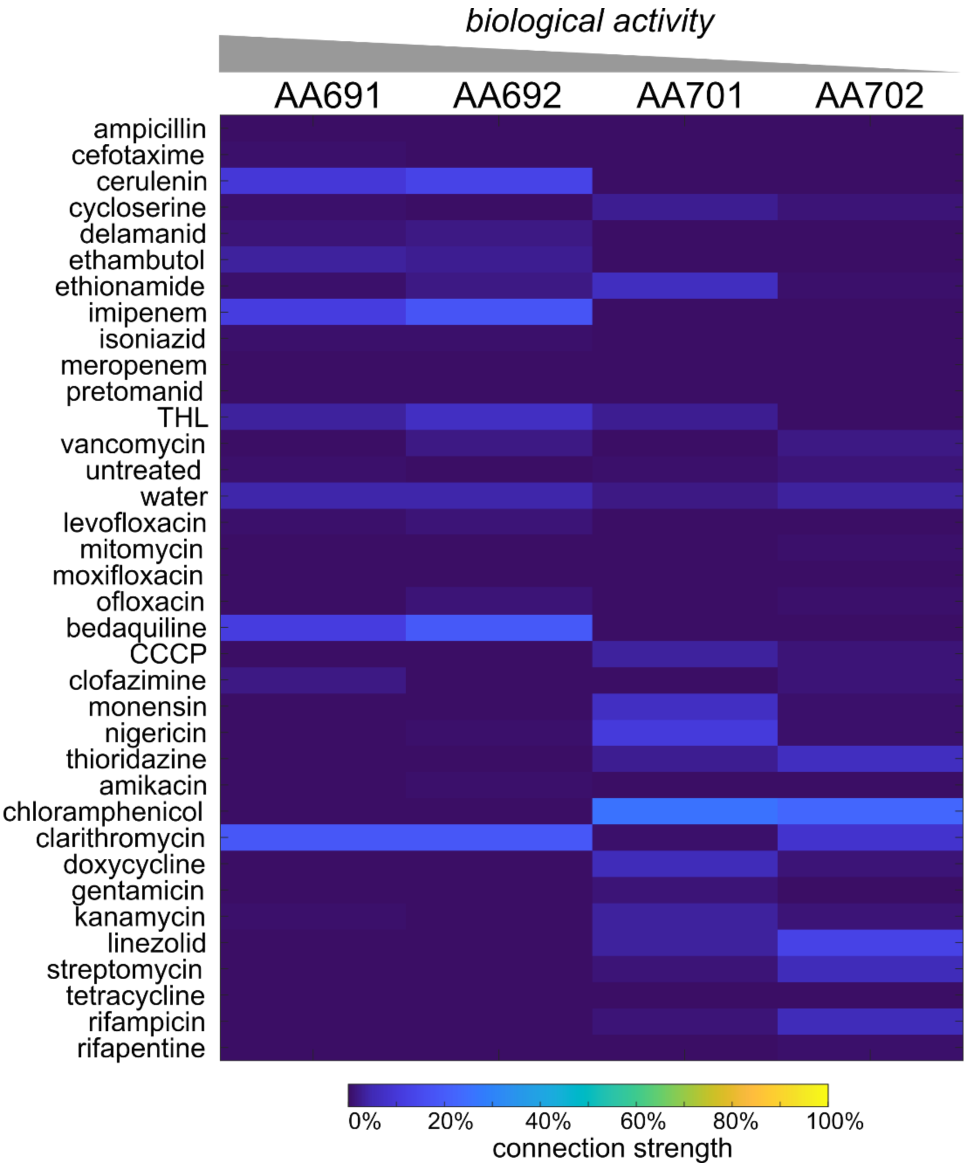


**Figure S7. AA691 and AA692 at low dose cause morphological changes in *Mtb* that are similar cell wall synthesis inhibitors and to the translation inhibitor clarithromycin.** *Mtb* were incubated with 50 μM compound (low dose) and stained for cellular membranes and the chromosomal nucleoid. Stained *Mtb* were imaged and analyzed for 25 morphological features. The profile for each compound was applied onto the morphological space constructed using 34 compounds with known mechanisms of action. The resulting nearest neighbor frequency (connection strength) based individual compounds is highest among drugs that cause similar types of cellular change.

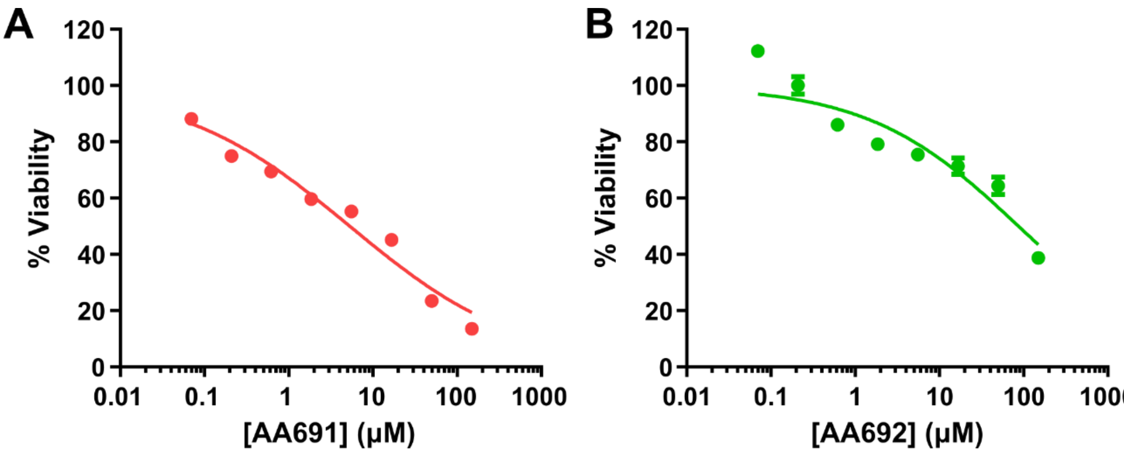


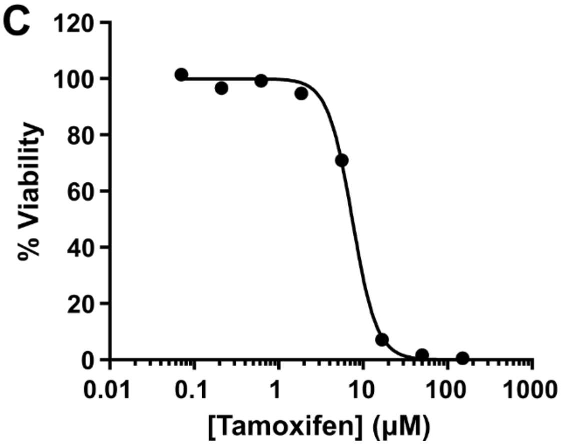


D

| Compound | HepG2 $CC_{50}$ (µM) | Mtb MIC (µM) | SI ($CC_{50}$/MIC) |
|---|---|---|---|
| AA691 | 5 ± 1 | 7 ± 4 | 0.7 ± 0.4 |
| AA692 | 89 ± 26 | 11 ± 4 | 7 ± 3 |
| Tamoxifen | 7.4 ± 0.2 | | |
| Isoniazid | 57,000[1] | 0.1 ± 0.04 | >570,000 |

**Figure S8. AA691 and AA692 are not highly selective for *Mtb*.** HepG2 cells were treated with A) AA691, B) AA692, or C) tamoxifen for 48 hours at 37 °C. Cell viability was measured using CellTiter-Glo (Promega). Data shown are the average ± S.D. of 3 technical replicates representative of 3 separate experiments. D) The selectivity index (SI) was calculated from the fit $CC_{50}$ and the MICs from Table 1. Data shown are the mean ± S.D. from 3 biological replicates. [1]From Elmorsy et al., Adverse effects of anti-tuberculosis drugs on HepG2 cell bioenergetics. *Hum Exp Toxicol*, 2017, 6, 616-625,

### SUPPLEMENTAL METHODS

### RESOURCE AVAILABILITY

***Materials availability***. Compounds synthesized for this study are available upon request. Strains generated for this study are available upon request.

***Data and code availability.*** Proteomics datasets from this study are deposited at ProteomeXchange. MorphEUS analysis data from this study are deposited at GitLab.

### EXPERIMENTAL MODEL AND SUBJECT DETAILS

***Bacterial strains and growth media***. *Mtb* H37Rv (BEI Resources NR-123) was used for ABPP and spontaneous resistant mutagenesis experiments. *Mtb* H37Rv harboring the integrating mLux plasmid (gift of Jeffery S. Cox), which expresses a codon-optimized bacterial *luxABCDE* operon for autoluminescence, was used for inhibitor screening, minimum inhibitor concentration (MIC) determination, and colony forming unit (CFU) enumeration. Autoluminescence has been previously validated as an indicator of *Mtb* viability (Zhang et al., 2012). Autoluminescent *M. smegmatis* (ATCC 700084) harboring the same integrating mLux plasmid; *E. coli* (strain MG1655) and *S. saprophyticus* (BAA-750, ATCC) were used for MIC determination.

For initial cultures frozen stocks of *Mtb* were thawed and used to inoculate Middlebrook 7H9 medium (BD) containing 10% (v/v) oleic acid-albumin-dextrose-catalase (OADC) supplement (BD), 0.5% glycerol, and 0.05% Tyloxapol (Sigma). Cells were then pelleted at 4000 x *g* for 10 minutes and washed once with modified Roisin’s medium (1 g/L $KH_2PO_4$, 2.5 g/L $Na_2HPO_4$, 5.9 g/L $NH_4Cl$, 2.0 g/L $K_2SO_4$, 1.0 g/L citric acid, 0.08 mg/L $ZnCl_2$, 0.4 mg/L $FeCl_3$-$6H_2O$, 0.02 mg/L $CuSO_4$, 0.02 mg/L $MnCl_2$-$4H_2O$, 0.02 mg/L $Na_2B_4O_7$-$10H_2O$, 0.02 mg/L $(NH_4)_6Mo_7O_{24}$-$4H_2O$, 0.5 mM $CaCl_2$, 0.5 mM $MgCl_2$, 0.5% glycerol, 0.5 mg/L biotin, 0.05% Tyloxapol, pH 6.6), and then resuspended in modified Roisin's medium. Acidic pH was achieved by buffering the medium to pH 5.0. In the acidic pH model, *Mtb* was cultured in pH 6.6, pelleted at 4000 x *g* for 10 minutes, washed once with pH 5.0 buffered modified Roisin's medium and incubated at pH 5.0 for 3 days as an adaptation step before further experimentation. *Mtb* cultures were incubated with shaking at 110 rpm or without shaking if in 96-well plates. *M. smegmatis* was inoculated from frozen stocks and cultured in modified Roisin's medium. *E. coli* and *S. saprophyticus* were similarly cultured in LB medium. All bacteria were cultured at 37 °C.

**METHOD DETAILS**

***Minimum inhibitory concentration (MIC) determination.*** Autoluminescent *Mtb* was subjected to the phenotypic screening protocol except compound concentrations varied from 100 µM to 0.2 µM. Autoluminescent *M. smegmatis* was subjected to the same protocol except incubation time was 10 hours (~3 doubling times) and the signal was measured with 400 ms integration time using a FilterMax F5 (Molecular Devices). *E. coli and S. saprophyticus* were cultured by an analogous procedure except that the incubation time was adjusted to ~3 doubling times for the respective bacterium (1.5 hours and 2 hours respectively). For *M. smegmatis* viability was measured by adding 100 µL BacTiter-Glo (Promega) to each well and incubating at 22 °C for 2 minutes before measuring luminescence as above. The linear range for each bacterium was determined by calibrating luminescence signal against $OD_{600}$. $MIC_{90}$ was determined by fitting the percent inhibition (versus DMSO vehicle-treated control) as a function of compound concentration to the Gompertz equation in GraphPad Prism 8. An estimated $MIC_{90}$ by visual inspection was determined by the same protocol, but in clear round-bottom 96-well plates. For this method the lowest concentration of compound at which no visible growth was observed was reported as the $MIC_{90}$.

***Compound toxicity.*** Synthesis of AA691-(1,4) and AA692-(1,4) and toxicity experiments were performed by Pharmaron. HepG2 cells were initially cultured in Dulbecco’s Modified Eagle’s Medium supplemented with 10% FBS (DMEM/FBS), 1x penicillin-streptomycin mixture and 1x non-essential amino acids. Medium was aspirated, 3 mL trypsin/EDTA solution was added, and cells were incubated at 37 °C for approximately 2 minutes or until the cells were detached and floated. Trypsin/EDTA was inactivated by adding DMEM/FBS. Cells were then centrifuged at 200 x *g* for 10 minutes. The supernatant was aspirated carefully and the cell pellet was re-suspended in DMEM/FBS. The cell density was adjusted to 8 x $10^4$ cells/mL and each well of a 96-well plate (Cellware) was seeded with 100 µL cell suspension. The medium was aspirated and 100 µL DMEM/FBS with compound or DMSO vehicle was added to the wells. Cells were incubated in a humidified, 37 °C, 5% $CO_2$ atmosphere for 48 hours. Subsequently, 50 µL pre-mixed Cell-Titer Glo (Promega) was added to each well and the plates were incubated at 22 °C for 10 minutes. Luminescence was measured on an Infinite M200 (Tecan). The percent signal was calculated by dividing the luminescent signal from a compound treated well by a vehicle treated well and multiplying by 100. The $CC_{50}$ was calculated by plotting the percent signal vs. compound concentration and fitting to the equation: Percent signal = Min + (Max - Min)/(1 + 10^((Log ($IC_{50}$) – Log (Concentration)) x (Hill Slope))) using GraphPad Prism.

***Compound stability.*** Compounds at a final concentration of 10 µM were incubated in modified Roisin's medium at pH 5.0 or pH 6.6 for 0, 1, 2, or 3 weeks at 37 °C. Samples were analyzed by liquid chromatography mass spectrometry (University of Illinois at Urbana-Champaign Mass Spectrometry Lab). Briefly, acetonitrile and formic acid were added to samples to final concentrations of 50% (v/v) and 0.1% (v/v), respectively. The MS analysis was conducted on an LTQ XL Orbitrap mass spectrometer (Thermo Fisher Scientific) operated in positive mode at a resolution of 30000. The spray voltage was 5.0 kV and the capillary temperature was 275 °C. The capillary voltage was 23 V. The m/z peaks corresponding to $[M + H]^+$, $[M + Na]^+$, and $[M + K]^+$, where M is a compound of interest, were calculated and manually assigned with the aid of ChemDraw (PerkinElmer). The ion count from each assigned m/z peak was used to monitor relative compound concentration over time.

***Enumeration of colony forming units (CFU)***. *Mtb* was cultured in 96-well plates as above for MIC determination in either pH 6.6 or pH 5.0 modified Roisin's medium. Separate plates were incubated for 0, 7, 14, or 21 days in a humidified environment at 37 °C and 5% $CO_2$. Where noted, hypoxia was achieved using a type A Bio-Bag environmental chamber (BD) according to the manufacturer's instructions. The chamber is reported to reach <1% oxygen within 2 hours and a resazurin indicator remained colorless within the Bio-Bag, confirming that the chamber remained hypoxic during the course of the experiment. At each time point, 10-fold serial dilutions ($1 \times 10^{-1}$, $1 \times 10^{-2}$, $1 \times 10^{-3}$, $1 \times 10^{-4}$ or $1 \times 10^{-2}$, $1 \times 10^{-3}$, $1 \times 10^{-4}$, $1 \times 10^{-5}$) from each well were plated on 7H11 Middlebrook agar with 10% OADC and 0.5% glycerol. CFU were enumerated after 3-4 weeks incubation at 37 °C, and 5% $CO_2$.

***Selection for resistant mutants.*** Roisin's solid medium was generated by adding 10 g/L bacterial agar (BD) to modified Roisin's medium. Compounds were added to the agar to a final concentration of 10x MIC. The final concentration of DMSO was 0.5% (v/v) for all plates. *Mtb* H37Rv was grown to an $OD_{600}$ 0.6-0.8 in modified Roisin's medium and an estimated 2 x $10^7$ and 2 x $10^8$ CFU were plated based on the assumption that an $OD_{600}$ 1 is approximately 3 x $10^8$ CFU/mL. Plates were incubated for 5-6 weeks in a humidified environment at 37 °C, 5% $CO_2$. The number of cells plated in each experiment was confirmed by plating 10-fold serial dilutions on Roisin's solid medium containing DMSO vehicle control.

***ABPP-SILAC for competitive ABPP at pH 5.0 and for detection of the active serine hydrolase proteome.*** Competitive ABPP-SILAC at pH 5.0 was performed as described for target identification by ABPP-SILAC but with the following changes: After culture growth to an $OD_{600}$ ~1 in light or heavy modified Roisin's medium at pH 6.6, cells were pelleted, washed twice with modified Roisin's medium at pH 5.0 and incubated in respective light or heavy modified Roisin's medium at pH 5.0 for 3 days. Light cultures were incubated with 13 μM AA691, AA692, AA701 or AA702 based on the approximate $MIC_{90}$ for AA692 (**Table 1**).

To identify the total active serine hydrolase proteome, ABPP-SILAC was performed as described above but with the following change: After light and heavy *Mtb* lysates were generated without compound treatment, heavy lysates were treated with 4 μM FP-biotin (gift of Dr. Eranthie Weerapana) (Liu et al., 1999) and light lysates were treated with 0.4% (v/v) DMSO vehicle control. We identified Rv0458 in all 15 instrumental replicates, it is probably not a serine hydrolase because (1) it is highly homologous to NADP-dependent oxidoreductases, which do not rely on serine-mediated catalysis, and (2) three enzymes (Rv3368c, Rv0927c, Rv2766c) that were detected in 12 or 13 of 15 replicates are also predicted oxidoreductases, but were not detected in the experiment comparing ABP treatment to vehicle control, suggesting that oxidoreductases were common contaminants in our biotin-streptavidin affinity enrichment.

***ABPP-SILAC mass spectrometry sample preparation and data analysis.*** Biotinylated proteins were enriched by incubating combined light/heavy lysates with 100 µL streptavidin agarose beads (Sigma-Aldrich) with gentle shaking at 25 °C for 2 hours. After removing the supernatant, the beads were washed three times with 0.25% SDS in 1 mL PBS, once with 1 mL PBS, and once with 1 mL deionized water for a total of five washes. The beads were then resuspended in 500 µL 6 M urea in PBS and treated with 25 µL 200 mM dithiothreitol in water for 15 min at 65 °C. The beads were then treated with 25 µL 400 mM iodoacetamide in water for 30 min at 37 °C. Samples were diluted with 950 µL PBS to stop the reaction, beads were pelleted at 1400 x *g* for 3 minutes, and the supernatant was aspirated. On-bead protease digestion was performed using 2 mg sequence-grade trypsin (Promega) in 2 M urea and 2 mM $CaCl_2$ in PBS for 12-14 hours at 37 °C. Peptides released by digestion were acidified with 5% formic acid and stored at -20 °C prior to analysis. Digested peptides were analyzed as described previously (Cognetta et al., 2015). Peptides were loaded onto a biphasic (strong cation exchange/reverse phase) capillary column and analyzed by 2D LC-MS/MS on a LTQ-

Orbitrap (Thermo Scientific). The ProLuCID algorithm was used to search spectra against a *Mtb* H37Rv reverse-concatenated nonredundant (gene-centric) FASTA database that was assembled from the UniProt database. SILAC ratios were quantified using in-house CIMAGE software (Weerapana et al., 2010).

For further analysis and comparison, SILAC ratios were converted to percent inhibition values. Note that all inhibition ratios >20 were reported as 20 and thus the maximum percent inhibition was 95. For the purposes of this study, negative percent inhibition (SILAC ratios >1) was treated numerically as zero inhibition. Under replicating conditions (pH 6.6), serine hydrolases that were inhibited 13% more by AA692 than AA702 (half a standard deviation greater than the mean of all inhibition values measured) were considered prioritized targets (**Table S5**). Using an analogous cutoff but under non-replicating conditions (pH 5.0), serine hydrolases that were inhibited 36% more by AA692 than AA702 were considered prioritized targets.

***Enzyme purification.*** The plasmid pET-15b_6His_Ag85A (BEI, NR-13292) encodes for Ag85A (Rv3804c), hereafter referred to as FbpA. This plasmid was transformed into *E. coli* BL21(DE3) cells and selected on LB plates containing 100 µg/mL carbenicillin (LB/carb100). A single carbenicillin resistant colony was grown in LB/carb100 overnight. This culture was used to inoculate 1 L of LB/carb100 and grown at 37 °C until $OD_{600}$ 0.6-0.8 was reached. Expression of FbpA was induced by 500 µM IPTG and the culture was grown at 18 °C for 16 hours. After induction, all purification steps were performed at 4 °C. Cells were centrifuged at 5000 x *g* for 20 min and the supernatant was removed. Cell pellets were resuspended in 30 mL lysis buffer (20 mM Tris, 200 mM NaCl, 1 mM DTT, 0.2 mM EDTA, 10 mM imidazole, 10% glycerol, pH 7.4) and sonicated with 5 s on/off for 10 min total processing time. The lysate was passed through a 0.45 μm syringe filter before loading onto a nickel affinity column (HisTrap FF 5mL, GE Healthcare). The column was washed with 5 column volumes of binding buffer (Buffer A: 50 mM Tris, 1 mM DTT, 10% glycerol, pH 7.4). Bound FbpA eluted at ~100 mM imidazole over a 0-50% gradient of elution buffer (Buffer A with 1 M imidazole) over 20 column volumes and purity was confirmed by SDS-PAGE.

*Mtb* fatty acid synthase Fas (Rv2524c) was purified as reported (Baron et al., 2018) from *E. coli* BL21(DE3) harboring the plasmids pMT100-Strep-Flag-*fas1* and pACYCDuet-Ara-*acpS* except that the cell lysate was loaded directly onto a 5 mL Strep-Trap HP column (GE Healthcare) without ammonium sulfate precipitation. Briefly, cells were grown at 37 °C in LB broth containing 100 μg/ml ampicillin and 17 μg/ml chloramphenicol to $OD_{600}$ 0.3-0.4. Expression of AcpS was induced by addition of 0.2% arabinose and the culture was grown to $OD_{600}$ 0.6-0.8. The expression of Fas was induced by addition of 0.5 mM IPTG at 15 °C for 20 hrs. The cells were harvested and resuspended in buffer A (100mM potassium phosphate pH 7.2, 150 mM KCl, 1 mM TCEP, 1 mM EDTA). After loading onto the Strep-Trap HP column, the column was washed with buffer A and eluted with buffer A containing 2.5 mM desthiobiotin. The thioesterase TesA (Rv2928) and the monoacylglycerol lipase Rv0183 were expressed and purified as reported (Cotes et al., 2007; Nguyen et al., 2018b). Briefly, for TesA, *E. coli* T7 Iq pLysS cells (New England Biolabs) harboring the plasmid pDEST14-TesA were grown in Terrific Broth to $OD_{600}$ 0.6–1.0 induced by with 0.5 mM IPTG overnight at 17 °C. Following cell harvesting and lysis by sonication in lysis buffer [50 mM Tris–HCl (pH 8), 300 mM NaCl, 10 mM imidazole, 0.25 mg/mL lysozyme], the supernatant was loaded onto HisTrap 5 mL (GE Healthcare). The protein was washed with buffer A [20 mM Tris–HCl (pH 8), 150 mM NaCl] containing 50 mM imidazole and eluted with buffer A containing 250 mM imidazole. Rv0183 was purified similarly, except *E. coli* Rosetta pLysS cells harboring pDEST14-His-Rv0183 were induced with 1 mM IPTG at 25 °C overnight and the lysis buffer was 50 mM Tris/HCl (pH 8.0), 150 mM NaCl, 1 mM EDTA, 0.1% Triton X-100, 0.25 mg/ml lysozyme.

***In vitro competitive ABPP assay.*** Purified proteins [Fas in 100 mM potassium phosphate buffer, pH 7.4; TesA or FbpA in 20 mM Tris-HCl, pH 7.4, 150 mM NaCl, 0.5% (w/v) Triton X-100] were incubated at 0.1 µM with various concentrations of compound for 2 hours at 22°C. For TesA the catalytically inactive mutant S104A (Nguyen et al., 2018b) was also included as a negative control. Appropriate mutants were not readily available for FbpA and Fas. For these enzymes, heat-treated samples (10 min at 93 °C) were included as a negative control. Subsequently, 5 µM TAMRA-FP (gift of Dr. Micah Niphakis) (Patricelli et al., 2001) was added and incubated for 1 hour at 22 °C. The final DMSO concentration was 2.5% (v/v). The reactions were quenched with the addition of 5x Laemmli sample buffer, separated by SDS-PAGE (10% for Fas; 15% for TesA, FbpA), and imaged with a Sapphire Biomolecular Imager (Azure) at 520 nm excitation wavelength and 575 nm emission wavelength. All image processing and analysis were performed using Image Studio Lite (LI-COR). Dose-response curves were fitted in GraphPad Prism8.

***Inhibition assays on recombinant TesA and Rv0183.*** The lipase-inhibitor preincubation method was used to test the direct inhibition of TesA or Rv0183 in presence of inhibitors as described previously (Nguyen et al., 2018a; Point et al., 2012). Briefly, an aliquot of each enzyme was pre-incubated at 25 °C with each inhibitor (50 µM stock solution in DMSO) at various inhibitor molar excess ($x_I$) ranging from 0.25 to 300 relative to 1 mol of enzyme. Preincubation with inhibitors was performed in the presence of 0.5% (*w/v*) Triton X-100 for TesA and 3 mM sodium taurodeoxycholate for Rv0183. A sample was collected after 30 min incubation and the residual enzyme activity was measured. The variation in the residual enzyme activity allowed determination of the inhibitor molar excess which reduced the activity to 50% of its initial value ($x_{I50}$). In each case, control experiments were performed in the absence of inhibitor. The respective enzymatic activity of TesA was assessed using the *para*-nitrophenyl (*p*NP) ester release assay with *p*NP valerate (*p*NP-C5) as substrate (Nguyen et al., 2018b). Rv0183 residual activity was determined potentiometrically using monoolein as substrate. Dose-response curves were fitted in Kaleidagraph 4.2 (Synergy Software).

**In silico *molecular docking experiments*.** Autodock Vina(Trott and Olson, 2009) was used as previously reported (Nguyen et al., 2017; Seeliger and de Groot, 2010) to generate the putative binding modes of the various inhibitors into the active site of TesA and the Rv0183. The PyMOL Molecular Graphics System (version 1.4, Schrödinger, LLC) was used as working environment with an in-house version of the AutoDock/Vina PyMOL plugin (Seeliger and de Groot, 2010). The X-ray crystallographic structure of TesA in complex with the $CyC_{17}$ inhibitor (PDB entry code: 6FVJ) and Rv0183 (PDB entry code: 6EIC) were used as receptors (Aschauer et al., 2018; Nguyen et al., 2018b). Docking runs were performed after replacing the catalytic serine (*i.e.*, Ser104 in TesA and Ser110 in Rv0183) by a glycine residue to enable the ligand (*i.e.*, the inhibitor) to adopt a suitable position corresponding to the pre-bound intermediate before the nucleophilic attack in the enzyme active site. The box size used for the various receptors was chosen to fit the whole enzyme's active site cleft and allowed non-constructive binding positions. The three-dimensional structures of the aforementioned compounds were constructed using Chem3D Ultra 11.0 software, and their geometry was refined using the Avogadro open-source molecular builder and visualization tool (version 1.2.0. http://avogadro.cc/).

### QUANTIFICATION AND STATISTICAL ANALYSIS

Statistical details of experiments, including number of independent experiments and/or replicates used for analysis; data reported; and statistical tests used, are provided in the figure legends. In general significance was determined as $p < 0.05$ (values for specific comparisons are also noted in the figure legends). Statistical analyses were performed with GraphPad Prism 8.

### SUPPLEMENTAL ITEMS

**Table S1**. Summary of ABPP-SILAC experiments

**Table S2.** Peptides detected in ABPP-SILAC experiments

**Table S3.** Active serine hydrolase proteome

**Table S4.** Prioritized serine hydrolase targets of AA692
